# Spatial Profiling Reveals Equivalence-Derived Molecular Signatures of Brain Mimicry and Adaptation in Breast Cancer Brain Metastases

**DOI:** 10.1101/2025.01.13.631781

**Authors:** Maxine Umeh Garcia, Christine Yiwen Yeh, Bryanna Godfrey, Pablo Nunez Perez, Giuseppe Barisano, Sushama Varma, Saman Ahmadian, Angus Toland, Monica Granucci, Thy Trinh, Hannes Vogel, Robert West, Michael Angelo, Lu Tian, Sylvia K. Plevritis, Melanie Hayden Gephart

**Affiliations:** Department of Neurosurgery, Stanford University School of Medicine, Stanford, CA 94305, USA; Department of Biomedical Data Science, Stanford University School of Medicine, Stanford, CA 94305, USA; Department of Pathology, Stanford University School of Medicine, Stanford, CA 94305, USA; Department of Neuropathology, Stanford University School of Medicine, Stanford, CA 94305, USA; Department of Radiology, Stanford University School of Medicine, Stanford, CA, 94305, USA; Department of Genetics, Stanford University School of Medicine, Stanford, CA, 94305, USA; Department of Medicine, Stanford University School of Medicine, Stanford, CA, 94305, USA

## Abstract

Brain metastases (BrMets), common for advanced-stage breast cancer patients, are associated with poor median survival and accompanied by severe neurologic decline. Halting the progression of breast cancer brain metastases (BCBMs) may require modulation of the tumor microenvironment (TME), yet little is known about the impact of the primary breast TME on brain tropism, or how, once there, metastatic breast cancer cells coexist with brain-resident cells (e.g., neurons and glia). Traditionally, studies in this space have focused on differential expression analysis, overlooking potential insights gained from investigating genes with equivalent expression between groups. This is particularly crucial in distant metastasis, where tumor cells may co-opt the transcriptional programs of the host organ (e.g., brain) to facilitate successful seeding and outgrowth. Prior to our work, no computational framework existed to determine biologically-relevant equivalent gene expression. To resolve molecular mechanisms of BCBM enabled by metastatic cancer cells and/or resident brain cells, we leveraged Nanostring GeoMx to perform spatially-resolved transcriptomic profiling on 235 patient-derived tissue cores from BCBM (including adjacent normal brain), primary invasive breast cancers, and normal (non-cancer) brain; analyzing 18,677 RNAs in 450 areas of interest (AOIs). We introduce the “Equivalent Expression Index” a highly specific and accurate algorithm that identifies statistically significant “Equivalently-Expressed Genes”. This method facilitated the identification of molecular remodeling and mimicry genes within tissue-specific TMEs. By integrating differential expression analysis with the Equivalent Expression Index, we discovered multiple novel gene signatures associated with BCBM and primary tumor brain-metastatic potential. We demonstrate that the Equivalent Expression Index is a powerful tool to uncover shared gene expression programs representing the adaptation of metastatic cells and brain-resident cells to the BCBM microenvironment.

## INTRODUCTION

Metastatic brain tumors, most frequently originating from lung, breast, and skin cancers, often represent highly-evolved, therapy-resistant tumors, occurring in ∼30% of patients.^1^ As treatments for cancer improve, patients survive longer, and neuroimaging techniques advance, the incidence and prevalence of brain metastasis increases.^2–4^ Overall, 10-15% of women with metastatic breast cancer develop brain metastases (BrMets), and rates as high as 30% (HER2+) and 50% (triple negative) are observed.^5^ The diagnosis of breast cancer brain metastasis (BCBM) is rapidly fatal, carrying a median survival time of 10 months, and is often accompanied by severe neurologic decline.^6^

The ability of metastatic cells to interact with organ resident cell types and remodel the host microenvironment determines metastatic capacity. This is particularly important in BrMets where the local environment is distinctly divergent from the primary breast tumor microenvironment. However, how metastatic cells adapt to their unique host environments, how the microenvironment is changed in the presence of cancer cells, and spatial resolution of the genes/pathways exploited by cellular subtypes to enhance adaptability, remain difficult to identify. While numerous studies have profiled differences between primary tumors and BrMets using differential expression analyses, and identified differentially expressed genes (DEGs), even DEGs shared between BrMets from different primary tumors types^7^, here we focus on spatially resolved, cell subtype-specific DEGs and Equivalently-Expressed Genes (EEGs) within the tumor microenvironment. EEGs may be particularly important in the context of distant metastasis, where metastatic tumor cells may exploit the transcriptional programs of the host organ (e.g., brain) to facilitate successful seeding and outgrowth. Specifically, metastatic cells may adapt to resemble the expression profiles of the distal organs they are colonizing, which would not be identified by differential expression. Similarly, the brain microenvironment may adapt to the presence of cancer cells – critically influencing tumor formation and disease progression.

To understand the BCBM microenvironment, we proposed an unbiased, spatially-resolved, quantitative approach to characterize biologically-relevant equivalent expression patterns in BCBM cell subtypes through use of the Equivalent Expression Index. The Equivalent Expression Index is a novel algorithm that identifies highly accurate, specific, and statistically significant Equivalently-Expressed Genes (EEGs) across cell types of interest. We explored how cancer and brain cells may employ shared transcriptional signals to effectively coexist in the BCBM tumor microenvironment (TME); and tested the hypothesis that metastatic breast cancer upregulated genes relevant to the brain microenvironment. We acquired spatial transcriptomic data from BCBM (including adjacent normal brain), primary invasive breast cancers, and normal (non-cancer) brain. We examined the expression of over 18,000 RNAs in 450 areas of interest (AOIs) enriched for malignant tumor cells, brain-resident cells (e.g., neurons, glia), or immune cells in patient-derived formalin-fixed, paraffin-embedded (FFPE) tissue sections, using the Nanostring GeoMx Digital Spatial Profiler (DSP).^8^

Leveraging the combination of differential expression analysis and the Equivalent Expression Index, we discovered multiple novel gene signatures associated with the clinical course of patients with BCBM. We demonstrated that upregulation of brain-associated EEGs by metastatic cancer cells (MIBS-9 signature) was associated with prolonged patient survival. Conversely, upregulation of cancer-associated EEGs by adjacent normal brain-resident cells (ARCS-81 signature) was associated with decreased survival. Moreover, we identified a brain-specific gene expression signature in primary breast tumors associated with the development of brain metastases (BACE-45). Taken together, our data underscore the power and broad utility of spatially-resolved coupling of differential expression analysis and the Equivalent Expression Index to elucidate novel genes reflective of brain tropism, microenvironmental support of BCBM, and TME adaptation.

## RESULTS

### Digital spatial transcriptomic profiling of breast tumors, brain metastases, and normal brain

We leveraged a combination of genome-wide transcriptome analysis and high-resolution imaging to spatially map the microenvironment of BCBM. We constructed a custom formalin-fixed, paraffin-embedded (FFPE) tissue microarray (TMA); which consisted of 235 tissue cores from 149 unique patients treated at Stanford Health Center between 2008-2019 (**Figure 1A**). Tumor sections were obtained from BCBM (n = 79) and primary invasive breast cancer (IBC) (n = 46). Of the IBC tissues, 24 were from patients who did not develop BrMets (“Non-Progressor-Primary”; confirmed by brain MRI), and 22 were from patients who subsequently developed BrMets (“Progressor-Primary”) within ∼10 years of initial breast cancer diagnosis. Sections were obtained from adjacent normal brain surrounding metastases (n = 48) and normal brain in patients without cancer (n = 23, temporal lobectomy tissue from epileptic patients). (**Supplemental Table 1**). Spatial transcriptomics data were collected using the Nanostring GeoMx Digital Spatial Profiler (DSP) platform^8^ depicted in **Figure 1B**, **1C**. Using data obtained from DSP, we then leveraged a combination of differential expression analysis and the Equivalent Expression Index (described later) using Boolean logic-based set analysis, to identify novel, biologically-relevant gene expression signatures associated with patient clinical outcomes (**Figure 1D**).

**Figure 1.**
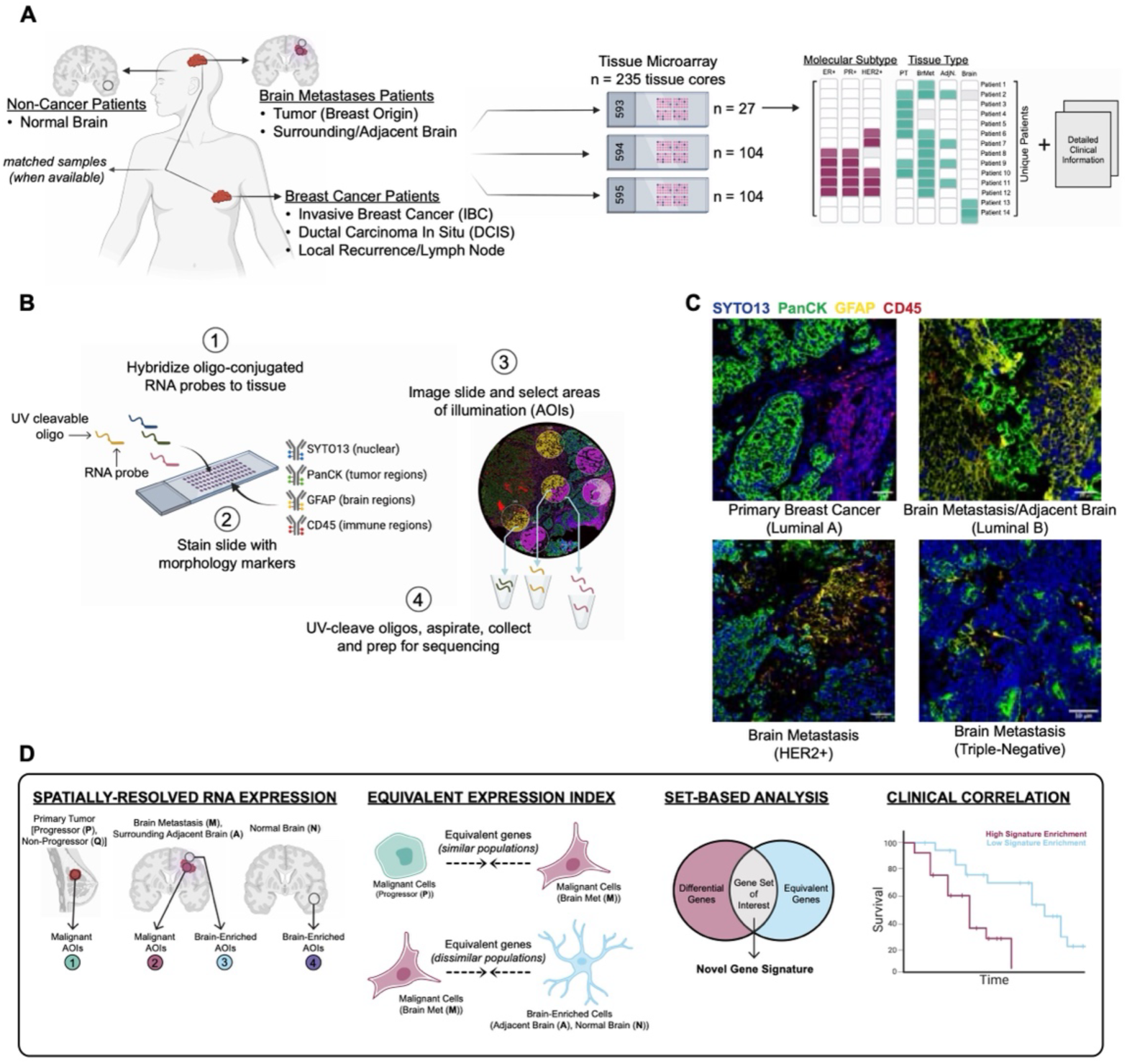
Digital spatial transcriptomic profiling of breast tumors, brain metastases, and normal brain. **(A)** Schematic of study cohort. Created with BioRender.com. Tissue was collected from breast cancer patients (n = 46), breast cancer brain metastases patients (n = 79) including surrounding brain tissue (n = 48), and non-cancer patients (n = 23), and organized across three FFPE tissue microarray blocks (n = 235 cores) with associated clinical data for each unique patient. **(B)** Schematic of digital spatial profiling (DSP) workflow with next generation sequencing readout. Created with BioRender.com. Whole transcriptome RNA probes/oligos are hybridized to tissue, tissue is stained with immunofluorescence (IF) markers to identify regions of interest, AOIs are selected and oligos are cleaved by UV light, oligos are collected and prepped for sequencing. **(C)** Representative IF-stained images of tissue regions with morphological markers SYTO13 (nuclear), PanCK (epithelial/tumor cells), GFAP (brain cells), and CD45 (immune cells). The scale bar is 10µm. **(D)** Schematic of study workflow. Created with BioRender.com.

We profiled a total of 473 AOIs across multiple tissue types and subtypes of breast cancer (**Figure 2A, 2B**), which were subsequently reduced to 450 AOIs following quality control filtering (**Supplemental Table 2**). We assigned AOIs to one of 9 cell types, based on immunofluorescence (IF) staining: Epithelial, Immune-Enriched, Malignant, Mixed0 [Malignant-Immune], Mixed1 [Malignant-Brain], Mixed2 [Brain-Immune], Mixed3 [Malignant-Brain-Immune], Brain-Enriched, Stroma-Enriched; and one of 6 tissue types, based on pathologist annotations: Normal Brain, DCIS, Local Recurrence/Lymph Node, BrMet, Other (Control Tissue), Primary Breast (IBC). Unbiased hierarchical clustering, based on pairwise correlation coefficients, revealed that cell types, and to a lesser extent tissue type and molecular subtype, affected the similarity between AOIs (**Figure 2C, 2D, Supplemental Figure 1A**). Following dimensionality reduction, AOIs clustered into three broad cell type groups: Immune-Enriched, Malignant, and Brain-Enriched (**Figure 2E**). AOIs containing a mix of cell types clustered according to the dominant cell type present in the AOI (**Supplemental Figure 1B, 1C**). Annotating AOIs by surface area, estimated nuclei count, assay slide, or collection plate revealed no relevant clusters, indicating that RNA profiles were not driven by these factors (**Supplemental Figure 1D-G**). Expression of known immune-, brain-, and tumor-associated genes confirmed cell type-specific gene expression across our broad (**Figure 2F**) and extended (**Supplemental Figure 1H**) cell type annotations. As expected, *PTPRC* (CD45) and *HLA-DRA*, expressed by antigen-presenting cells^9^, were significantly elevated in Immune-Enriched AOIs. Glial fibrillary acidic protein (*GFAP*)^10^ and myelin basic protein (*MBP*)^11^, highly expressed in the brain, were significantly and specifically elevated in Brain-Enriched AOIs; and *KRT8*^12^ and *EpCAM*^13^, highly expressed in tumor cells, were significantly elevated in Malignant AOIs. AOI annotations underwent additional, multistep validation (Methods and **Supplemental Figure 2**).

**Figure 2.**
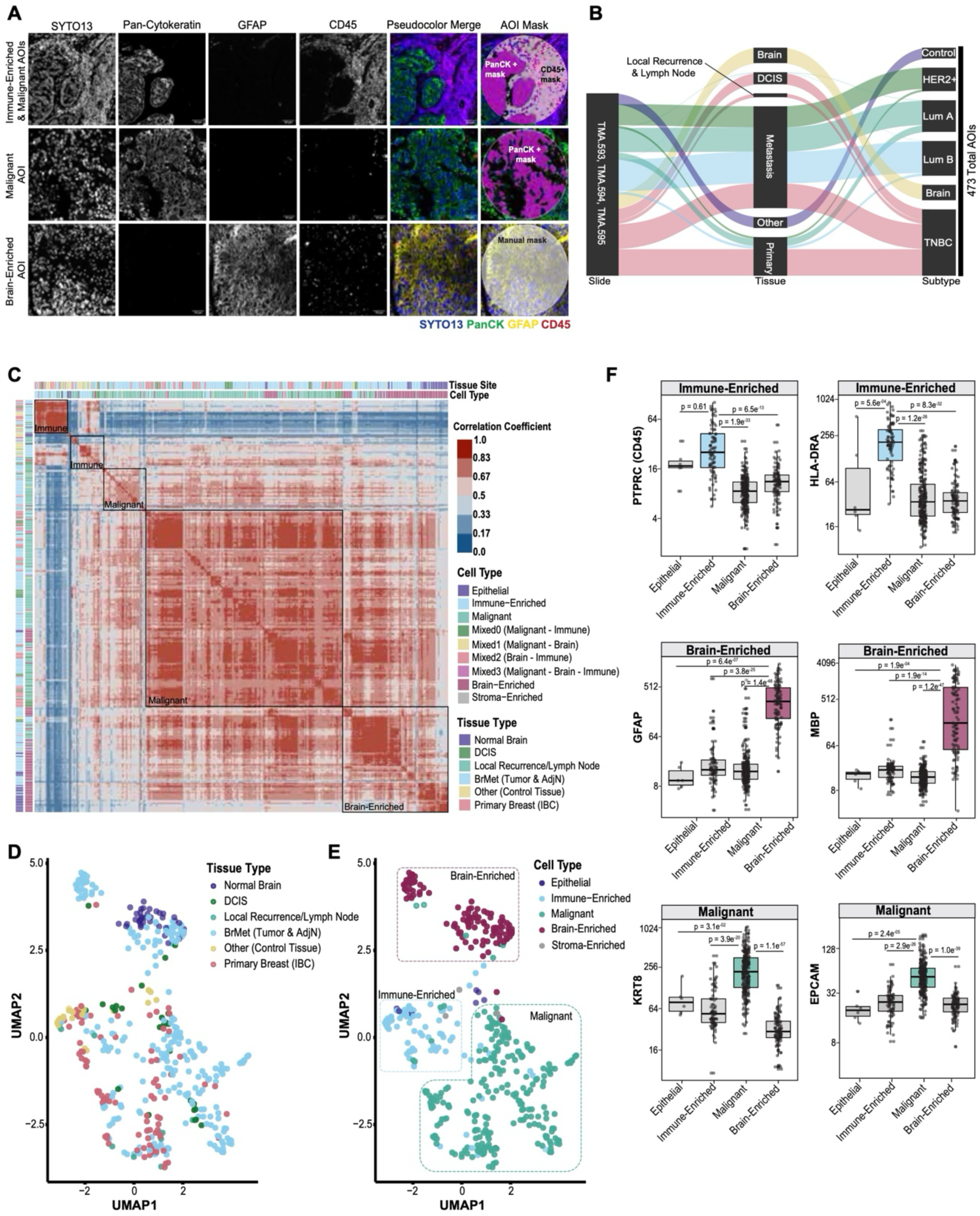
Cell type and tissue type influence the expression profile of each ROI. **(A)** Representative example of Immune-Enriched, Malignant, and Brain-Enriched AOIs. Individual IF channels, pseudocolor merged IF image, and AOI masked images are shown. AOIs selected were on average 200µm circles. **(B)** Sankey diagram visually summarizing the experimental design showing slides, tissue types, and molecular subtypes AOIs profiled (n = 473). **(C)** Correlation matrix heatmap showing the pairwise correlation coefficients (Pearson) between AOIs (n=450) using Q3 normalized counts (n = 9575 genes). Heatmap colored from blue to red according to correlation coefficient score 0.0 to 1.0. Tissue type and cell type are indicated by colored bars. The five largest clusters (hclust method) are boxed and named according to the predominate AOI type (Immune-Enriched, Malignant, Brain-Enriched). **(D-E)** UMAP clustering of AOIs annotated by **(D)** tissue type and **(E)** cell type. **(F)** Boxplots of Q3 normalized gene counts by AOI type for selected immune-, brain-, and tumor-associated genes. n = 6, 83, 248, and 111 AOIs in Epithelial, Immune-Enriched, Malignant, and Brain-Enriched, respectively. For boxplots in **F**, middle line denotes the median, box edges indicate the 25th and 75th percentiles, and whiskers extend to the most extreme points that do not exceed ±1.5 times the interquartile range (IQR). P values are based on nonparametric test (Kruskal–Wallis) followed by Dunn test for pairwise comparisons. DCIS: Ductal carcinoma in situ. TNBC: Triple-Negative breast cancer. UMAP: Uniform Manifold Approximation and Projection.

### Differential gene expression patterns in the brain metastases microenvironment

Consistent with traditional methods that utilize differential expression analyses, we initially investigated transcriptional changes within both the metastatic tumor cells and the brain microenvironment. We focused on understanding how primary breast cancer cells differed from BCBM, and how the presence of tumor cells influenced transcriptional signals in adjacent normal brain regions. We identified a gene signature which captured transcriptional differences in Malignant AOIs between primary tumors of patients who progressed to brain metastases [Progressor-Primary (**P**); (n=34)] and BCBM [Metastasis (**M**); (n=154)]. We likewise compared Brain-Enriched AOIs from brain adjacent to BCBM [Adjacent Normal (**A**); (n=73)] to patients without cancer [Normal Brain (**N**); (n=38)] (**Figure 3A**). From these analyses we derived BCBM-309 (where the last component of the signature name denotes the number of genes included). BCBM-309 consisted of 309 DEGs: 45 genes up-regulated in Metastasis (**Figure 3B, 3C**) compared to primary tumor, and 264 genes up-regulated in Adjacent Normal (**Figure 3D, 3E**) compared to non-cancer brain. Gene set enrichment analysis confirmed the association between BCBM-309 [Enriched in BCBM (**M** + **A**)] and diverse process including stemness (*CTBP2*, *PDCD2*), neurodegeneration (*DDX5*, *DUSP6*, *TFRC*, *STXBP5L*), MTORC1 signaling (*CANX*, *SCD*, *TFRC*), and metastasis (*KRT18*, *HMGA1*,*TMED9*, *TMED3*, *SCGB2A2*) (**Figure 3F**). We also investigated transcriptional changes in immune cell populations (Immune-Enriched AOIs) between Progressor-Primary (**P**) (n=10) and Metastasis (**M**) (n=28) (**Supplemental Figure 3A, 3B**). Top DEGs in Metastasis Immune-Enriched AOIs included *MUC1*, *AGRN*, and *ST3GAL1*; as well as *LAMA5*, and *ALCAM*, shown to regulate T-cell mediated immune responses and endothelial cell composition, including blood brain barrier maintenance, respectively.^14–18^ Moreover, markers of an “exhausted environment”^19^ in breast tumors (specifically exhausted T-cells: *CD8B*, *LAG3*, *CAMK1D*, and *DPF3*) were increased in Metastasis (**M**) Immune-Enriched AOIs, supporting the idea that the metastatic TME may be more immunologically inert than the primary tumor TME.^20,21^

**Figure 3.**
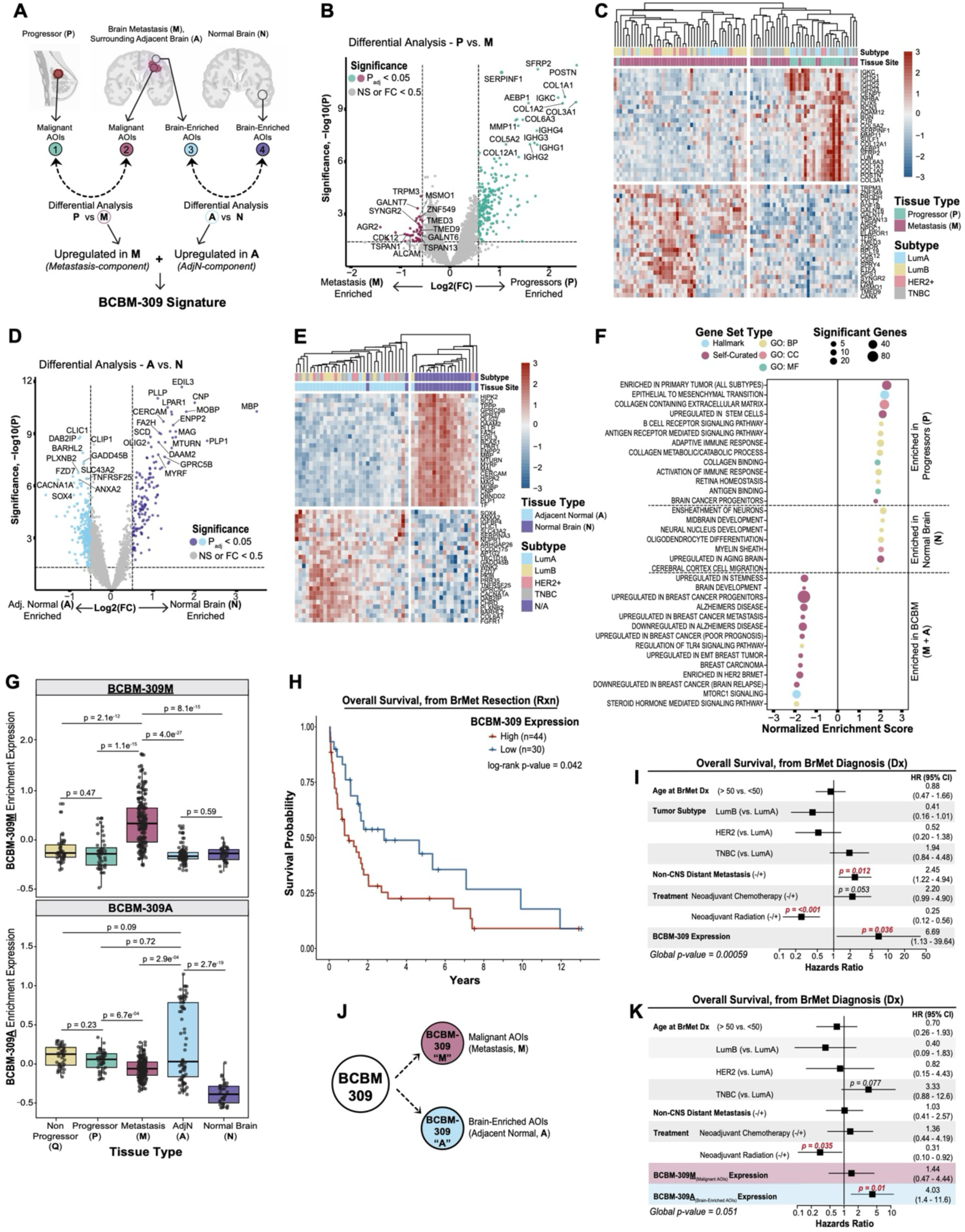
Spatially-resolved gene expression patterns in the brain metastases microenvironment. **(A)** Schematic of BCBM-309 signature derivation. Created with BioRender.com. Genes upregulated in Metastasis compared to Progressor-Primary were combined with genes upregulated in Adjacent Normal compared to Normal Brain. **(B)** Volcano plot of differentially expressed genes between Progressor-Primary (n = 34) and Metastasis (n =154) Malignant AOIs. **(C)** Heatmap of top 25 differentially expressed genes in Progressor-Primary and Metastasis Malignant AOIs. **(D)** Volcano plot of differentially expressed genes between Adjacent Normal (n = 73) and Normal Brain (n = 38) Brain-Enriched AOIs. **(E)** Heatmap of top 25 differentially expressed genes in Adjacent Normal and Normal Brain Brain-Enriched AOIs. **(F)** Gene set enrichment analysis of genes upregulated in each group: Progressor, Normal Brain, BCBM – Metastasis & Adjacent Normal. Gene set type and number of significant (p < 0.05) genes are denoted. **(G)** Boxplots of BCBM-309M and BCBM-309A enrichment expression by tissue type. n = 43, 44, 182, 73, and 38 AOIs in Non-Progressor, Progressor, Metastasis, Adjacent Normal, Normal Brain; respectively. For boxplots in **G**, middle line denotes the median, box edges indicate the 25th and 75th percentiles, and whiskers extend to the most extreme points that do not exceed ±1.5 times the interquartile range (IQR). P values are based on nonparametric test (Kruskal–Wallis) followed by Dunn test for pairwise comparisons. **(H)** Kaplan-Meier curves of overall survival (from date of brain metastases resection) as a function of BCBM-309 expression. Patients were segregated into two groups (High, n = 44; Low, n = 30) based on the enrichment expression of the BCBM-309 signature in Metastasis and Adjacent Normal AOIs. The log-rank p-value was derived from comparing discretized predictors (high versus low BCBM-309 expression). **(I)** Multivariable Cox proportional hazards regression analysis assessing the impact of clinical covariates and the expression of BCBM-309 on overall survival (from date of brain metastasis diagnosis). **(J)** Schematic of BCBM-309 signature deconvolution into two components: BCBM-309M (Malignant AOIs from Metastasis) and BCBM-309A (Brain-Enriched AOIs from Adjacent Normal). **(K)** Multivariable Cox proportional hazards regression analysis assessing the impact of clinical covariates and the expression of BCBM-309M and BCBM-309A on overall survival (from date of brain metastasis diagnosis). In **B** and **D**, color coding refers to the AOI types in **A**. In **B** and **D**, p-values were calculated using a mixed-effects model using chi-square (χ2) tests (two-sided) with p < 0.05 considered as significant. Genes with p > 0.05 and/or Log_2_(Fold Change) < 0.5 are considered not significant. In **C** and **E**, tissue type and molecular subtype are indicated by colored bars. Heatmaps colored from blue to red according to Log_2_(FC) value of –3 to 3. In **I** and **K**, for each covariate, the hazard ratio (HR) and its 95% confidence interval (CI), denoted by bars, are presented. In **I** and **K**, p-values for each covariate were calculated using the Wald statistic. p < 0.05 are significant and denoted in bold red text. p-values near significance are also noted. Global p-values were calculated using the log-rank statistic. FC: fold change. GO: Gene ontology. BP: biological process. CC: cellular component. MF: molecular function. Rxn: Resection. Dx: Diagnosis.

### Reprogramming of brain-resident cell types by the presence of metastatic cancer correlates with survival outcomes

We then computed the enrichment of the two components of the BCBM-309 signature: the metastasis-component (BCBM-309**<u>M</u>**, referring to the 45 genes upregulated in Metastasis) and the adjacent normal-component (BCBM-309**<u>A</u>**, referring to the 264 genes upregulated in adjacent normal brain) across all AOIs (n = 380) from five tissue types [Non-Progressor-Primary (**Q**), Progressor-Primary (**P**), Metastasis (**M**), Adjacent Normal (**A**), and Normal Brain (**N**)] (**Figure 3G**). Since many of the genes in BCBM-309 have not previously been described in BrMets, we cross-referenced the signature with publicly available patient-matched BCBM data.^22^ Overall expression of BCBM-309**<u>M</u>** and BCBM-309**<u>A</u>** was significantly increased in BrMets relative to matched primary breast tumors (p=4.8e^−05^, p=0.0069) (**Supplemental Figure 3C, 3D**). Additionally, scRNA-Seq data^23^ demonstrated that BCBM-309**<u>M</u>** was specific to metastatic tumor cells, BCBM-309**<u>A</u>** was most enriched in macrophages (likely including brain-resident microglia), and BCBM-309 overall was an accurate integration of both cell types (**Supplemental Figure 3E**).

We next explored the association between BCBM-309 and patient overall survival. Patients had significantly decreased survival time associated with higher BCBM-309 expression alone (**Figure 3H**, log-rank p=0.042), as well as when accounting for additional clinical covariates (**Figure 3I**, Cox regression p=0.036, global p=0.00059). Again, portioning BCBM-309 into the Malignant (Metastasis, **M**) and Brain-Enriched (Adjacent Normal, **A**) AOIs, we tested the correlation with overall survival outcomes (**Figure 3J**). This revealed that although metastatic tumor cell expression was correlated with shorter survival outcomes, it was largely expression from brain-resident cells that drove the increased hazard of death (**Figure 3K**, Cox regression p=0.01, global p=0.05). This finding held true when we examined BCBM-309A expression as an independent predictor of overall survival (**Supplemental Figure 3F**, log-rank p=0.049), as well as when controlling for clinical covariates (**Supplemental Figure 3G**, Cox regression p = 0.031, global p=0.017). These findings demonstrate that using traditional differential expression analysis we can identify DEGs correlated with overall survival outcomes in BCBM, particularly when considering adjacent normal brain. However, by extending our analyses beyond differential expression, and leveraging the importance of Equivalently-Expressed Genes (EEGs), we were able to further deconvolute these transcriptional programs.

### A computational framework for identifying Equivalently-Expressed Genes (EEGs)

Non-differentially expressed genes are not by default equivalent. Therefore, to quantitatively and unbiasedly identify spatially-resolved signatures of transcriptional programs shared between cell subtypes in the BCBM TME (e.g., between tumor and brain-resident cells), we implemented the novel and broadly applicable Equivalent Expression Index. In contrast to the traditional approach of differential expression analysis, as used to derive the BCBM-309 signature (**Figure 4A**), the Equivalent Expression Index identified statistically significant Equivalently-Expressed Genes (EEGs) across AOIs with high accuracy and specificity (**Figure 4B**), regardless of tissue type or cell subtype.

**Figure 4.**
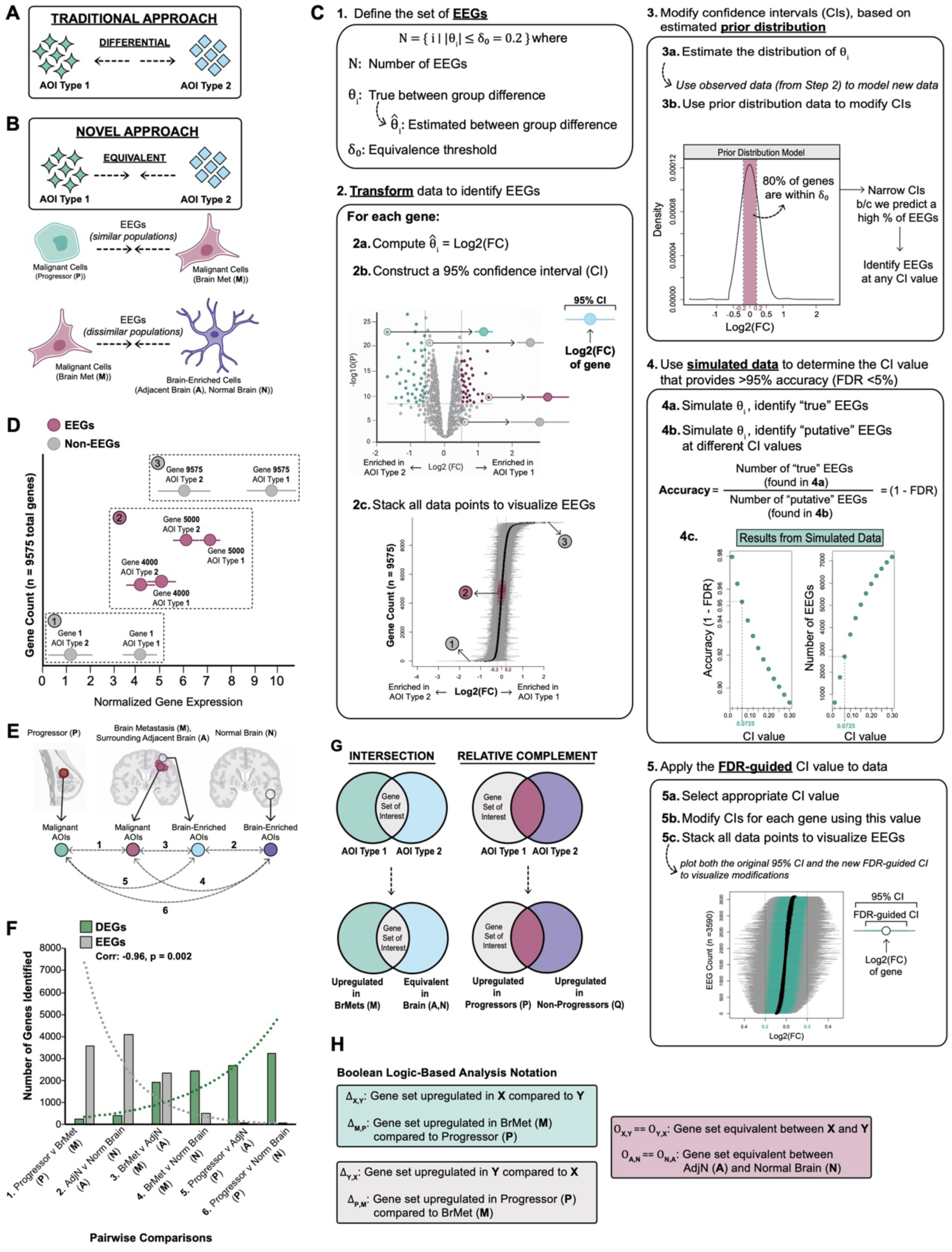
The Equivalent Expression Index enables identification of Equivalently-Expressed Genes (EEGs). **(A)** Schematic of traditional differential expression analyses which identify genes of interest based on divergence between AOI types. **(B)** Schematic of novel equivalent expression analyses which identify genes of interest based on equivalence between AOI types. Schematic depicting that equivalent expression can be assessed between similar and/or dissimilar AOI types. **(C)** Detailed schematic of Equivalent Expression Index workflow: define set of EEGs, transform data to identify EEGs, modify confidence intervals based on prior distribution, simulate data to determine confidence interval thresholds with accuracy, apply FDR-guided confidence intervals to data. **(D)** Schematic of EEGs versus non-EEGs. Plot of normalized gene expression (X-axis) depicting four genes of interest (corresponding to three regions in the plot in 4D.2c). Y-axis corresponds to gene number label. Genes in grey represent non-EEGs with significant separation in normalized gene expression between AOI types. Genes in maroon represent EEGs with little to no separation in normalized gene expression between AOI types. **(E)** Schematic of the six pairwise equivalent expression analyses carried out between AOI types: (1) Malignant AOIs from Progressor-Primary (P) versus Malignant AOIs from Metastasis (M), (2) Brain-Enriched AOIs from Adjacent Normal (A) versus Brain-Enriched AOIs from Normal Brain (N), (3) Malignant AOIs from Metastasis (M) versus Brain-Enriched AOIs from Adjacent Normal (A), (4) Malignant AOIs from Metastasis (M) versus Normal Brain (N), (5) Malignant AOIs from Progressor-Primary (P) versus Brain-Enriched AOIs from Adjacent Normal (A), (6) Malignant AOIs from Progressor-Primary (P) versus Brain-Enriched AOIs from Normal Brain (N). Created with BioRender.com. **(F)** Barplot depicting the total number of differentially expressed genes (DEGs) versus equivalently expressed genes (EEGs) in the six pairwise comparisons outlined in **E**. An exponential trendline is fitted to the data. Pearson correlation coefficient and p-value are provided. The p-value is calculated using a two-sided t-test. **(G)** Venn diagram illustrating representative Boolean Logic-based set analyses to identify gene sets of interest (e.g., intersection, relative complements), including representative examples used in this study (bottom). **(H)** Equations showing the generalizable notation developed for this study, including examples of notation applied to AOI types. EEGs: Equivalently Expressed Genes. DEGs: Differentially Expressed Genes. FC: fold change. CI: confidence interval. FDR: false discovery rate.

Analytically, we defined EEGs as genes for which the absolute value of the expression difference between two AOI types fell within a predefined interval known as the Equivalence threshold (δ_0_) (**Figure 4C.1**). For these studies we set δ_0_ to (–0.2, 0.2). Genes that did not meet these requirements were considered non-EEGs (and not necessarily DEGs). To identify EEGs, for each gene in our dataset (n=9,575), we first computed the estimated between group difference (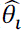), equal to the Log_2_(Fold Change), between the two AOI types being compared. We then constructed a 95% confidence interval (CI) around each value and stacked these transformed data points to visualize EEGs (**Figure 4D.2a-c**), being genes whose Log_2_(Fold Change) and 95% CI are contained within the (–0.2, 0.2) interval. Conceptually, EEGs exhibited little to no difference in expression in the two AOI types being compared (**Figure 4D**, region 2), while non-EEGs exhibited larger differences in gene expression between AOI types (**Figure 4D**, regions 1 and 3). As shown in **Figure 4D.2c** (region 2), only a small number of EEGs are identified due to this highly stringent approach. Moreover, the false discovery rate (FDR) of this approach is unknown, and likely very low (<1%), resulting in a high number of false negatives (i.e., genes that are EEGs but not identified). Since our goal was to take a discovery-based approach to identify EEGs, we reduced stringency by modifying the CIs around each gene using a prior distribution model based on observed data (**Figure 3D.3a-b, Supplemental Figure 4A, 4B**). For example, if the prior distribution estimated that 80% of genes within our dataset had a Log_2_(FC) between –0.2 and 0.2, we narrow the CIs around each gene such that more genes fell within the Equivalence threshold (–0.2, 0.2) and were identified as EEGs. This resulted in substantially more EEGs (**Supplemental Table 3**). However, because this is a less stringent method, we needed to account for the accuracy of this approach. To this end, we used simulated data to determine the CI value that provided a FDR of less than 5%, corresponding to >95% accuracy (**Figure 4D.4a-b**). To accomplish this, we simulated *θ_i_* for each gene (true Log_2_(FC) between groups) and identified “true” EEGs. We then simulated 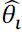 (estimated Log_2_(FC) between groups) for each gene and identified “putative” EEGs at various CI values. To calculate accuracy, we divided the number of “true” EEGs by the number of “putative” EEGs at each CI value. As expected, we observed an inverse correlation between accuracy and the number of EEGs (**Figure 4D.4c**). Given the tradeoff between accuracy and the number of EEGs, we selected the highest CI value which produced >95% accuracy. Lastly, we applied the selected CI value to our data, modifying each gene’s CI such that we identified a high number of EEGs while accounting for accuracy (**Figure 4D.5a-c**). (For all analyses in this study, we selected the CI value that corresponded to FDR of <5%). Further details of the Equivalent Expression Index are described in Methods.

Using the Equivalent Expression Index, we identified statistically significant EEGs between each pairwise comparison of AOI types (**Figure 4E**, **Supplemental Table 3**). This analysis revealed that similar AOI types (e.g., Brain-Enriched AOIs from Adjacent Normal (**A**) and Normal Brain (**N**)) exhibited a high number of EEGs, while dissimilar AOI types (e.g., Malignant AOIs from Progressor-Primary (**P**) and Brain-Enriched AOIs from Normal Brain (**N**)) exhibited lower numbers of EEGs. The number of EEGs showed a strong inverse correlation with the number of DEGs between comparisons (**Figure 4F**, correlation coefficient=-0.96, p=0.002). We then leveraged the unique combination of our differential expression analyses and Equivalent Expression Index data, using Boolean logic-based set analysis (**Figure 4G**); including the development of a generalizable notation (**Figure 4H**), to describe how we derived the novel genes signatures introduced below. Collectively, this method, and associated validations (**Supplemental Figure 4C – 4E**), demonstrate that the Equivalent Expression Index can be used to identify EEGs with high accuracy and specificity, across similar and dissimilar tissues and cell subtypes. Development of this novel tool, while broadly applicable, allowed us to explore shared transcriptional signals within the BCBM microenvironment.

### The Equivalent Expression Index reveals transcriptional mimicry in the brain metastases microenvironment

Using a combination of differential expression analysis and the Equivalent Expression Index, we derived two novel gene signatures that explore BrMet-specific shared transcriptional signals. First, we postulated that in comparison to primary tumors, BCBM may be facilitated, in part, by the ability to upregulate genes relevant in the brain microenvironment, potentially enhancing the ability of metastatic cells to coexist in the brain (i.e., upregulated genes in BCBM that are equivalently expressed by brain-resident cells). Thus, we intersected DEGs upregulated in Metastasis (versus Progressor-Primary) (Δ_M,P_) with EEGs between Brain-Enriched AOIs (Adjacent Normal and Normal Brain) ($_A,N_) (**Figure 5A**). This identified 9 genes: *MSMO1, RPL38, SYNGR2, CANX, ZNF549, XYLT2, ELAPOR1, GALNT7, REEP5* – termed the MIBS-9 (**<u>M</u>**etastasis enr**I**ched in **<u>B</u>**rain-**<u>S</u>**hared – **<u>9</u>** gene) signature or “brain equivalence” signature. Genes in the MIBS-9 signature have been associated with cancer,^24–26^ but, intriguingly, also have recognized roles in the central nervous system and neurodegeneration.^27–32^ As expected, the overall expression of MIBS-9 was significantly enriched in Metastasis (**M**) AOIs compared to Progressor-Primary (**P**) (**Figure 5B**, p=1.7e^−12^), and when we assessed the expression of each of the 9 genes individually, we found no significant difference between Adjacent Normal (**A**) and Normal Brain (**N**) AOIs (**Supplemental Figure 5A**). Overlaying gene expression onto AOI images revealed, as expected, that MIBS-9 was highly enriched in Metastasis (**M**) AOIs (versus Progressor-Primary (**P**) AOIs), and equivalently expressed in Adjacent Normal (**A**) and Normal Brain (**N**) AOIs (**Figure 5C**), further confirming that this novel signature accurately reflects spatial gene expression patterns of distinct cell subtypes. Additionally, we found increased expression of MIBS-9 in HER2+ and Lum B Metastasis (**M**) AOIs, compared to Lum A and TNBC; with TNBC showing the lowest expression of all molecular subtypes (**Supplemental Figure 5B**, p≤2.6e^−06^). We observed no difference in the expression of MIBS-9 across molecular subtypes for Adjacent Normal (**A**) AOIs, suggesting that varying levels of MIBS-9 was specific to malignant cells within the BCBM TME. Due to the restricted resolution of brain-specific cell types (e.g., neurons, glia) in our Brain-Enriched AOIs (and lack of publicly available scRNA-Seq data that fully captures brain-resident cells in metastases), we turned to the GTex Portal^33^ and BrainRNASeq.org^34^ to investigate the expression of MIBS-9 across normal brain regions and brain-resident cell types, respectively. Although, MIBS-9 genes were variably expressed across brain anatomical sites, genes generally clustered into two groups: those expressed in the cerebellum and cortex (*REEP5*, *XYLT2*, *ZNF549*, *GALNT7*), and those expressed in the spinal cord, breast, and lung (*ELAPOR1*, *SYNGR2*, *RPL38*, *CANX*, *MSMO1*) (**Supplemental Figure 5C**). Analysis of brain cell type-specific data revealed high levels of MIBS-9 genes – *RPL38*, *CANX*, *MSMO1*, and *REEP5* – in brain-resident cells, specifically, oligodendrocytes, neurons, and microglia (**Figure 5D**, **Supplemental Figure 5D**). We then assessed the association between MIBS-9 and patient survival time. This revealed that high MIBS-9 expression was significantly associated with increased overall survival time (**Figure 5E**, log-rank p=0.00083, **Figure 5F**, Univariate Cox regression p=0.041, global p=0.003), suggesting that MIBS-9 genes may be functionally important.

**Figure 5.**
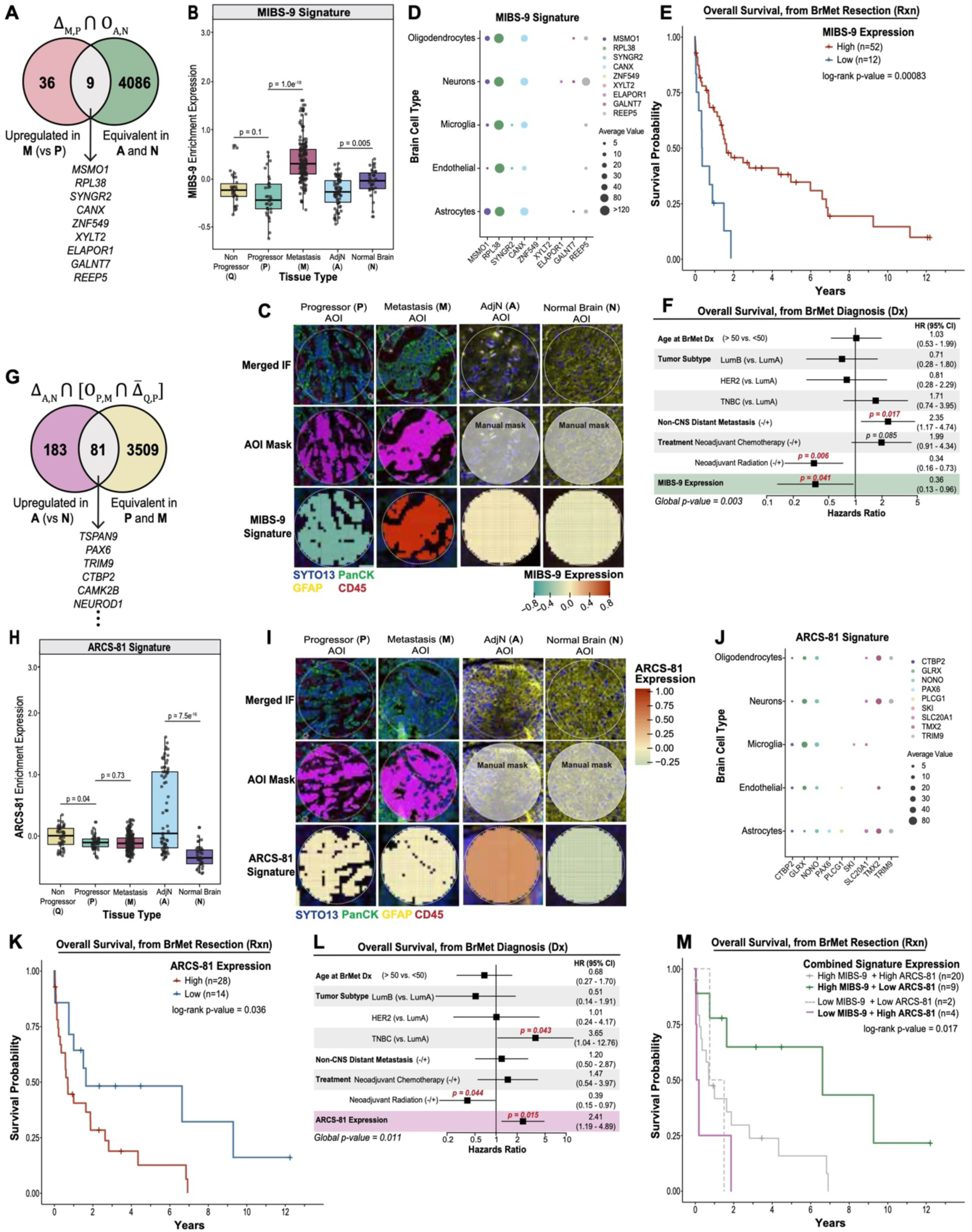
Transcriptional mimicry in the brain metastases microenvironment. **(A)** Venn diagram illustrating the set of genes that overlap between genes upregulated in Metastasis [compared to Progressor-Primary] Malignant AOIs and genes equivalent between Adjacent Normal and Normal Brain Brain-Enriched AOIs. Notation of set analysis is shown above Venn diagram. **(B)** Boxplot of MIBS-9 enrichment expression by tissue type. n = 43, 44, 182, 73, and 38 AOIs in Non-Progressor, Progressor, Metastasis, Adjacent Normal, Normal Brain; respectively. **(C)** Representative spatial omics overlay of MIBS-9 enrichment by tissue type. **(D)** Dot plot of expression values of MIBS-9 genes by brain cell type. **(E)** Kaplan-Meier curves of overall survival (from date of brain metastases resection) as a function of MIBS-9 expression. Patients were segregated into two groups (High, n = 52; Low, n = 12) based on the enrichment expression of the MIBS-9 signature in Metastasis AOIs. **(F)** Multivariable Cox proportional hazards regression analysis assessing the impact of clinical covariates and the expression MIBS-9 on overall survival (from date of brain metastasis diagnosis). **(G)** Venn diagram illustrating the set of genes that overlap between genes upregulated in Adjacent Normal [compared to Normal Brain] Brain-Enriched AOIs and genes equivalent between Progressor-Primary and Metastasis Malignant AOIs. Notation of set analysis is shown above Venn diagram. **(H)** Boxplot of ARCS-81 enrichment expression by tissue type. n = 43, 44, 182, 73, and 38 AOIs in Non-Progressor, Progressor, Metastasis, Adjacent Normal, Normal Brain; respectively. **(I)** Representative spatial omics overlay of ARCS-81 enrichment by tissue type. **(J)** Dot plot of expression values of ARCS-81 genes by brain cell type. **(K)** Kaplan-Meier curves of overall survival (from date of brain metastases resection) as a function of ARCS-81 expression. Patients were segregated into two groups (High, n = 28; Low, n = 14) based on the enrichment expression of the ARCS-81 signature in Brain-Enriched AOIs. **(L)** Multivariable Cox proportional hazards regression analysis assessing the impact of clinical covariates and the expression ARCS-81 on overall survival (from date of brain metastasis diagnosis). **(M)** Kaplan-Meier curves of overall survival (from date of brain metastases resection) as a function of the combination of MIBS-9 and ARCS-81 expression. Patients were segregated into four groups (High MIBS-9 + High ARCS-81, n = 20; High MIBS-9 + Low ARCS-81, n = 9; Low MIBS-9 + Low ARCS-81, n = 2; Low MIBS-9 + High ARCS-81, n = 4) based on the enrichment expression of the MIBS-9 and ARCS-81 signatures in Malignant and Brain-Enriched AOIs; respectively. For boxplots in **B** and **H**, middle line denotes the median, box edges indicate the 25th and 75th percentiles, and whiskers extend to the most extreme points that do not exceed ±1.5 times the interquartile range (IQR). P values are based on nonparametric test (Kruskal–Wallis) followed by Dunn test for pairwise comparisons. In **C** and **I**, pseudocolor merged IF image, AOI masked image, and signature enrichment overlay are shown. Signature enrichment overlays were generated using the SpatialOmicsOverlay R package (see Methods). In **D** and **J**, dot color and size represent the gene of interest and average expression value, respectively. In **E**, **K**, and **M**, the log-rank p-value was derived from comparing discretized predictors (high versus low expression of MIBS-9 (**E**), ARCS-81 (**K**), combination of MIBS-9 and ARCS-81 (**M**)). In **F** and **L**, for each covariate, the hazard ratio (HR) and its 95% confidence interval (CI), denoted by bars, are presented. In **F** and **L**, p-values for each covariate were calculated using the Wald statistic. p < 0.05 are significant and denoted in bold red text. p-values near significance are also noted. Global p-values were calculated using the log-rank statistic. Δ: upregulation. 0: equivalence. ∩: intersection. Rxn: resection. Dx: diagnosis.

Having identified a “brain equivalence” expression signature in metastatic tumor cells, we next sought to investigate whether the converse may be occurring; particularly, transcriptional changes in the Adjacent Normal that may support coexistence with metastatic cells. We postulated that genes upregulated in Adjacent Normal (versus non-cancer brain) and equivalent to BCBM may reflect the brain’s response to breast cancer exposure. We intersected DEGs upregulated in Adjacent Normal (versus Normal Brain) (Δ_A,N_) with EEGs between Malignant AOIs (Progressor-Primary and Metastasis) ($_P,M_) (**Figure 5G**). To focus on genes facilitating brain-specific metastasis, from our breast cancer EEG set ($_P,M_) we excluded any genes that were upregulated in Malignant AOIs of Non-Progressor-Primary (**Q**) (Δ_Q,P_), the breast cancer patients that did not progress to brain metastases. This produced an 81 gene signature termed **<u>A</u>**djacent normal en**<u>R</u>**iched in **<u>C</u>**ancer-**<u>S</u>**hared – **<u>81</u>** gene (ARCS-81). Notably, this novel gene signature contained brain-associated genes known to play a role in cancer (e.g., *NEUROD1*^35,36^, *TSPAN9*^37^*, CTBP2*^38,39^, *CAMK2*^40^). As expected, ARCS-81 expression was significantly higher in Adjacent Normal (**A**) AOIs versus Normal Brain (**N**) (p=7.8e^−11^); with near equal expression in AOIs from Progressor-Primary (**P**) and Metastasis (**M**) (median: –0.09, –010; respectively) (**Figure 5H**). This was further confirmed by overlaying ARCS-81 expression onto AOI images (**Figure 5I**). Expression of each of the 81 genes of interest followed the expected pattern of upregulation in Adjacent Normal (**A**) and equivalent expression in Malignant AOIs (**Supplemental Figure 5E**). Interestingly, in contrast to the MIBS-9 signature, which had the highest expression in HER2+ and LumB BrMets, ARCS-81 was lowest in HER2+ (**Supplemental Figure 5F**, p≤0.01), suggesting that these two signatures, while not mutually exclusive, may be specific to breast cancer molecular subtypes. Although, ARCS-81 genes were variably expressed across brain anatomical sites, genes generally clustered into two groups: those specifically expressed in the cerebellum (e.g., *TSPAN9*, *NEUROD1*, *PAX6*, *CAMKK2*), and those concurrently expressed in the cerebellum, breast, and lung (e.g., *NONO*, *CTBP2*, *TMX2*, *SLC20A1*, *PLCG1*) (**Supplemental Figure 5G**). As expected, given that the ARCS-81 signature was derived from Brain-Enriched AOIs, ARCS-81 genes were widely expressed across brain-resident cell types (**Figure 5J**) notably, neurons, astrocytes, and microglia (**Supplemental Figure 5H**). High ARCS-81 expression was significantly associated with decreased overall survival time (**Figure 5K**, log-rank p=0.036, **Figure 5L**, Univariate Cox regression p=0.015, global p=0.011) suggesting that ARCS-81 signature expression may be important in BCBM progression.

MIBS-9 and ARCS-81 were derived from metastatic tumor cells (**M**) and Adjacent Normal brain cells (**A**), the two components which together make up a large part of the BCBM TME. Thus, we also assessed the impact of combining both signatures on prediction of patient survival. Stratifying patients by the combination of high versus low MIBS-9 and ARCS-81 expression revealed that expression of high MIBS-9 (in metastatic cells) and low ARCS-81 (in Adjacent Normal regions) was associated with significantly longer overall survival times, while the combination of low MIBS-9 and high ARCS-81 was associated with significantly shorter survival time (**Figure 5M**, log-rank p=0.017). The inclusion of clinical covariates into this assessment supported the opposing impact of these signatures, and further emphasized the role of Adjacent Normal brain in survival correlations (**Supplemental Figure 5I,** Univariate Cox regression p=0.007, global p=0.045). Collectively, these results demonstrate that we can use the Equivalent Expression Index to identify EEGs present between diverse AOI (Malignant versus Brain-Enriched) and tissue (Progressor-Primary versus Metastasis) types; and that these novel, shared transcriptional programs are both biologically relevant and clinically significant.

### Progressor-specific primary tumor profiling reveals genes associated with brain-specific metastasis

Previous studies suggested that metastatic competence of tumor cells is present early in cancer development, long before metastases are clinically evident;^41^ and may explain why expression profiles of primary tumors can be used to predict the likelihood of distant metastases.^42^ To identify biologically-relevant gene signatures within primary breast tumors associated with the development of brain-specific distant metastases, we compared Malignant AOIs of Non-Progressor-Primary (**Q**) to Progressor-Primary (**P**) by differential expression analysis (**Figure 6A**). We identified 484 DEGs upregulated in Progressor-Primary compared to non-progressors (**Figure 6B**). Interestingly, the most upregulated gene was SF3B1, the most commonly mutated RNA splicing factor in cancer^43^, suggesting that aberrant RNA splicing, leading to widespread defects in RNA and protein production, may support cancer development and enhance metastatic potential. We validated the Progressor-upregulated gene set, termed **<u>P</u>**rogressor-**<u>S</u>**pecific **<u>S</u>**ignature – **<u>484</u>** gene (PSS-484), in three publicly available breast cancer datasets focused on pathological complete response (pCR, Wolf et al.^44^), risk and status of distant metastasis (Cheng et al.^45^), and time to distant metastasis (Wang et al.^46^). Overall expression of PSS-484 was higher in patients who did not achieve pCR (**Supplemental Figure 6A**, p=0.03), were at high risk for, and who developed, distant metastasis (**Supplemental Figure 6B**, p=1.8e^−07^, p=0.02); and had shorter time to distant metastasis development (**Supplemental Figure 6C**, log-rank p=0.0024); supporting the role of PSS-484 genes in the context of primary breast cancer metastatic potential. Gene set enrichment analysis revealed pathways known to be involved in metastatic competence, including EMT (*IGFBP2*, *THBS1*, *BGN*), angiogenesis (*VEGFA*, *LRPAP1*), negative regulation of intrinsic apoptotic signaling (*BCL2*, *HIF1A*, *XBP1*), endothelial cell chemotaxis (*HSPB1*, *FGFR1*), regulation of fibroblast migration (*AKT1*, *PTK2*), and RNA splicing (*SF3B1*, *DDX17*, *SF1*, *RALY*, *ESRP2*) (**Figure 6C**). PSS-484 also showed significant enrichment for cholesterol homeostasis (*ATF3*, *TP53INP1*, *CD9*), fatty acid metabolism (*ACSS1*, *LDHA*, *MDH1*, *ALDOA*), and brain-related genes (*CD63*, *SELENOP*, *COX6C*). We also investigated transcriptional differences in Immune-Enriched AOIs from Progressor-Primary (**P**) and Non-Progressor-Primary (**Q**) (**Supplemental Figure 6D, 6E**). Interestingly, *SCG5*, a secreted chaperone protein, with high expression in brain and BCBMs^47^, and involved in the regulation of adiposity^48^, was significantly upregulated in Progressor-Primary immune cell populations. Likewise, enrichment analysis of genes upregulated in Progressor-Primary (**P**) Immune-Enriched AOIs revealed processes associated with angiogenesis and EMT, similar to PSS-484 enrichment analysis; whereas Non-Progressor-Primary (**Q**) immune populations were enriched for gene sets related to immune response (*RNF168*, *HLA-A*, *HLA-B*, *TAP1)* (**Supplemental Figure 6F**). Overall, these data suggest an ability to predict brain tropism in primary cancer, while also pointing to a potential role for immune surveillance in the primary tumor that may contribute to immune tolerance of brain metastases.

**Figure 6.**
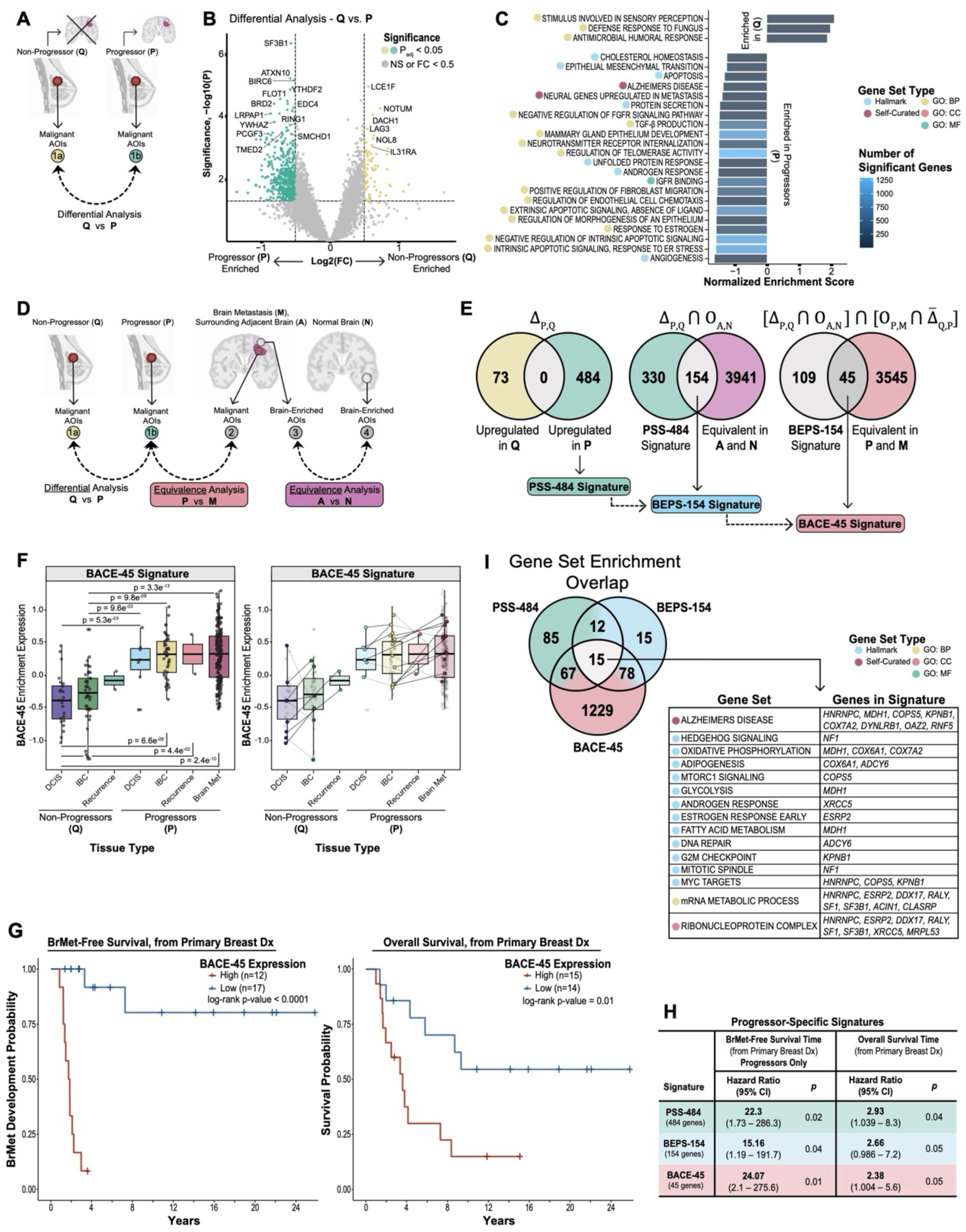
Progressor-Primary-Specific signature genes predict brain specific metastasis. **(A)** Schematic of Progressor-Primary-specific analysis. Created with BioRender.com. Differential gene expression analysis was performed between Non-Progressor-Primary and Progressor-Primary Malignant AOIs. **(B)** Volcano plot of differentially expressed genes between Non-Progressor-Primary (n = 29) and Progressor-Primary (n =34) Malignant AOIs. Color coding refers to the AOI types in **A**. P-values were calculated using a mixed-effects model using chi-square (χ2) tests (two-sided) with p < 0.05 considered as significant. Genes with p > 0.05 and/or Log_2_(Fold Change) < 0.5 are considered not significant. **(C)** Gene set enrichment analysis of genes upregulated in Non-Progressor-Primary compared to Progressor-Primary. Gene set type and number of significant (p < 0.05) genes are denoted. **(D)** Schematic of Progressor-Primary-specific signature derivations based on differential expression analysis. Created with BioRender.com. Non-Progressor-Primary versus Progressor-Primary Malignant AOIs, and equivalent expression analysis: Progressor-Primary and Metastasis Malignant AOIs, Adjacent Normal and Normal Brain Brain-Enriched AOIs. **(E)** Venn diagrams illustrating the derivation of Progressor-Primary-specific gene signatures. (1) Left Venn diagram shows the set of genes that overlap between genes upregulated in Non-Progressor-Primary and Progressor-Primary in Malignant AOIs (PSS-484 is derived from genes upregulated in Progressor-Primary [compared to Non-Progressor-Primary]). (2) Middle Venn diagram shows the set of genes that overlap between PSS-484 and genes equivalent between Adjacent Normal and Normal Brain Brain-Enriched AOIs (BEPS-154 is derived from overlapping genes). (3) Right Venn diagram shows the set of genes that overlap between BEPS-154 and genes equivalent between Progressor-Primary and Metastasis Malignant AOIs, after removing any genes upregulated in Non-Progressor-Primary (BACE-45 is derived from overlapping genes). Notation of set analyses is shown above Venn diagrams. (F) Boxplot of BACE-45 enrichment expression by tissue type over disease progression (*left*) and highlighting paired samples (*right*). In *right*, lines connect paired samples: solid lines indicate increased expression between pairs, dotted lines indicate decreased expression between pairs. Non-Progressor group: n = 24, 44, and 2 AOIs in DCIS, IBC, and Recurrence; respectively. Progressor group: n = 7, 45, 2, and 182 AOIs in DCIS, IBC, Recurrence, and Metastasis; respectively. For boxplots in F, middle line denotes the median, box edges indicate the 25th and 75th percentiles, and whiskers extend to the most extreme points that do not exceed ±1.5 times the interquartile range (IQR). P values are based on nonparametric test (Kruskal–Wallis) followed by Dunn test for pairwise comparisons. **(G)** Kaplan-Meier curves of brain metastasis-free survival (BMFS) (from date of primary breast cancer diagnosis) (*left*) and overall survival (OS) (from date of primary breast cancer diagnosis) (*right*) and as a function of BACE-45 expression. Patients were segregated into two groups for BMFS (High, n = 12; Low, n = 17) and OS (High, n = 15, Low, n = 14) based on the enrichment expression of the BACE-45 signature in Malignant AOIs from primary breast tumors. The log-rank p-value was derived from comparing discretized predictors (high versus low BACE-45 expression). **(H)** Summary of multivariable Cox proportional hazards regression analyses assessing the impact of clinical covariates (not shown, detailed in Supplemental Figure 7E-7F) and the expression PSS-484, BEPS-154, and BACE-45 on brain metastasis-free survival (BMFS) (from date of primary breast cancer diagnosis) and overall survival (OS) (from date of primary breast cancer diagnosis). In **H**, the hazard ratio (HR) for the gene signature of interest and its associated p-value are presented. The p-value was calculated using the Wald statistic with p < 0.05 considered as significant. **(I)** Venn diagram illustrating the gene sets that overlap between PSS-484, BEPS-154, and BACE-45 signatures based on gene set enrichment analyses. Table shows details of overlapping genes sets and associated genes. FC: fold change. DCIS: ductal carcinoma in situ. IBC: invasive breast cancer. GO: gene ontology. BP: biological process. CC: cellular component. MF: molecular function. Dx: diagnosis.

### The Equivalent Expression Index refines key features of the primary breast TME associated with brain metastasis

To further explore brain-related gene expression in primary breast cancer and its association with brain metastasis development, we examined genes that were upregulated in Progressor-Primary Malignant AOIs (DEGs between **Q** and **P**, (Δ_P,Q_)), equivalent in Brain-Enriched AOIs (EEGs between **A** and **N**, ($_A,N_)), and consistently maintained during the transition from primary to metastatic disease (EEGs between **P** and **M**, ($_P,M_)) (**Figure 6D**). To this end, we derived two additional gene signatures in a stepwise fashion to identify novel genes present in primary breast cancer, maintained in metastases, and reflective of the brain TME (**Figure 6E**). To begin, we intersected PSS-484 (Δ_P,Q_) with EEGs between Brain-Enriched AOIs (Adjacent Normal and Normal Brain) ($_A,N_). This identified 154 genes, termed **<u>B</u>**rain **<u>E</u>**quivalent **<u>P</u>**rogressor **<u>S</u>**pecific – **<u>154</u>** (BEPS-154), which included many cancer– and metastasis-associated genes such as BCL2,^49^ OTUB1,^50^ CD164/MUC24,^51,52^ and NF1.^53,54^ We then intersected the BEPS-154 signature (Δ_P,Q_ ∩ $_A,N_) with EEGs between Malignant AOIs (Progressor-Primary and Metastasis) ($_P,M_), excluding any genes upregulated in Non-Progressor-Primary compared to Progressor-Primary (Δ_Q,P_). This refined the gene set to 45 genes of interest termed **<u>B</u>**rain **<u>A</u>**nd **<u>C</u>**ancer **<u>E</u>**quivalence – **<u>45</u>** (BACE-45). As expected, the overall expression of the three signatures was significantly higher in Progressor-Primary (**P**) compared to Non-Progressor-Primary (**Q**) (**Supplemental Figure 6G**, p≤1.2e^−08^). Notably, these signatures were not only significantly upregulated in primary tumors of progressing patients, but their expression was also elevated over disease progression, with higher enrichment in invasive breast cancer (IBC) versus ductal carcinoma in situ (DCIS) – a precursor of breast cancer, and further elevated in the metastatic setting, as evidenced by increased expression in matched patient samples (**Figure 6F**, BACE-45 signature shown). Overlaying the three signatures onto AOIs confirmed enrichment in Progressor-Primary (**P**) AOIs and equivalence between Adjacent Normal (**A**) and Normal Brain (**N**) AOIs (**Supplemental Figure 6H**). We validated the BEPS-154 and BACE-45 signatures using the same publicly available datasets used for PSS-484, finding that expression of both signatures were higher in patients who did not achieve pCR (**Supplemental Figure 6I**, BEPS-154: p=0.021, BACE-45: p=0.081), had increased risk (BEPS-154: p=0.007, BACE-45: p=0.001) and development of distant metastasis (BEPS-154: p=0.11, BACE-45: p=0.041) (**Supplemental Figure 7A**), and reduced time to metastasis development (**Supplemental Figure 7B**, BEPS-154: log-rank p=0.01, BACE-45: log-rank p=0.041). We then explored the association of these signatures with both brain metastasis-free survival (BMFS) and overall survival (OS) in our cohort. This revealed that high PSS-484, BEPS-154, and BACE-45 were associated with decreased time to brain metastasis development and reduced overall survival time (**Figure 6G, Supplemental Figure 7C, 7D**, BMFS: log-rank p<0.0001, OS: log-rank p=0.01), even upon controlling for clinical covariates (**Supplemental Figure 7E, 7F**). Constraining this analysis to only patients that developed BrMets confirmed that higher expression of PSS-484, BEPS-154, and BACE-45 was significantly associated with shorter time to BrMet development (log-rank p=0.02) but was not associated with overall survival times (log-rank p=0.2) (**Supplemental Figure 7G**). A summary of these findings is depicted in **Figure 6H**. To identify key pathways across these signatures, we intersected gene set enrichment analysis data, revealing 15 gene sets present in all three signatures, containing a diverse array of biological processes, including those related to Alzheimer’s Disease (*MDH1*, *COPS5*, *KPNB1*, *OAZ2*), adipogenesis (*ADCY6*), and fatty acid metabolism/glycolysis (*MDH1*) (**Figure 6I**). Taken together, by coupling Progressor-Primary-upregulated DEGs with brain– and metastasis-specific EEGs, we identified novel genes of interest; highlighting molecular pathways in both the primary breast and normal brain that may be important for brain-specific metastatic potential.

## DISCUSSION

In this study, we combined genome-wide transcriptome analysis with high resolution imaging to acquire RNA expression data from spatially-resolved tissue regions (Areas of Interest, AOIs). Profiling of distinct AOIs from primary tumors, BrMets, and non-cancer brain suggests that the brain TME is remodeled to support a niche for BrMets, and that brain-tropic cues are present in the primary breast TME (**Figure 7**). We present the Equivalent Expression Index as a novel method to identify Equivalently-Expressed Genes (EEGs) between AOI types. We then leveraged a unique combination of differential and equivalence expression, using Boolean logic-based set analysis, to study shared gene expression patterns in both the primary and metastatic TMEs; identifying novel BCBM and primary tumor-specific gene signatures correlated with patient survival. Overall, our findings support use of the Equivalent Expression Index as a means to identify statistically-significant, biologically-relevant EEGs; uncovering shared gene expression signatures that may facilitate the adaptation of metastatic cells to the brain microenvironment.

**Figure 7.**
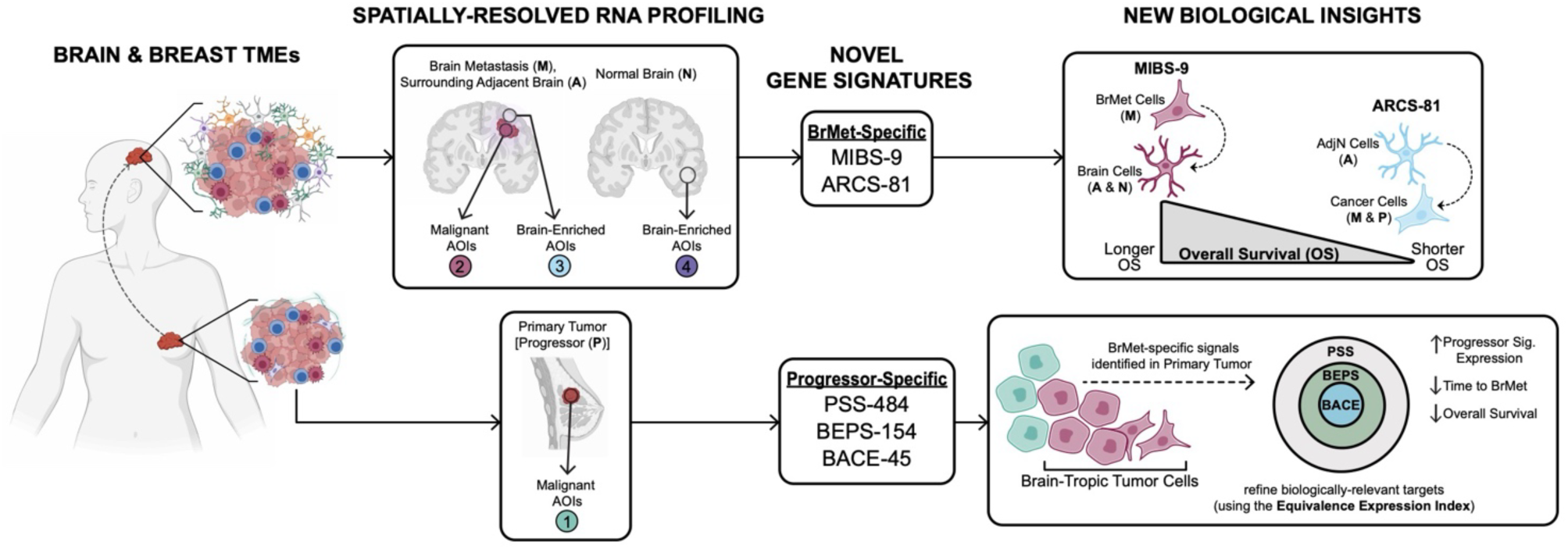
Summary schematic. A graphical summary depicting the major findings of this study. Created with BioRender.com. RNA spatial profiling was used to derive novel brain– and breast-specific gene signatures which were translated to novel biological insights.

It is not fully understood why only a subset of highly metastatic cancers have a proclivity for metastasis to the brain. Although the brain microenvironment is highly divergent from the primary TME, up to 40% of patients with metastatic breast, lung, and melanoma develop brain metastases, in contrast to the fewer than 1% of patients with prostate or pancreatic cancer. Brain metastasis from breast cancer typically occurs years after breast cancer diagnosis and frequently after diagnosis of metastatic disease, suggesting that cancer cells initially lack full competence to grow in the brain, and likely acquire these features due to selective microenvironment and/or treatment pressures.^7^ This adaptive ability may be encoded by gene regulatory programs that allow cancer cells to exploit aspects of the host microenvironment, or by the local microenvironment’s adaptation to the presence of cancer cells.^55^

Despite its significant therapeutic potential, there have been limited strategies for large-scale, unbiased investigations into how cancer cells adapt to the unique brain microenvironment, and how the microenvironment itself adjusts in response to the presence of cancer cells. We aimed to study TME interactions in BCBM within their native morphological context using GeoMx Digital Spatial Profiling (DSP). Spatial resolution was critically important to ensure that expression profiles were not affected by the inadvertent inclusion of other cell types (e.g., adjacent normal brain regions were not “contaminated” with tumor cells). DSP allows manual, user-defined selection of AOIs, including visual inspection during and post-experiment to confirm that regions are cell type specific. After validating the quality and reproducibility of DSP data, we captured the expression of 9,575 genes across 450 AOIs, making this the largest spatial profiling study of BCBM to date. Furthermore, the compatibility of DSP with FFPE tissue allowed us to profile cells in adjacent normal brain regions, which have largely been absent in scRNA-Seq studies, making the involvement of brain-resident cells in the pathology of BrMets poorly understood. Interestingly, we found that transcriptional signals from the adjacent normal brain are significantly associated with patient survival outcomes, highlighting the importance of these cell types.

We classified the BCBM TME into two interconnected components – Metastasis (M), which was enriched in malignant and immune cells and Adjacent Normal brain (A), which was enriched in brain-resident cells. Compared to primary breast tumors that ultimately develop brain metastases (Primary-Progressor (P)), the BrMet TME is characterized by markers of stemness, neurodegeneration, and immune cell exhaustion. This suggests that the BrMet TME supports outgrowth of metastatic cells, while immune responses against the tumor are suppressed, and may contribute to neurological damage. Spatially resolved, cell type-specific DSP data has considerable advantage over bulk RNA measurements, which tend to obscure less abundant brain-resident cell types. While the 45 BrMet-upregulated genes were also enriched in bulk RNA-Seq data of BCBM, only with our approach were we able to identify the important contribution of adjacent normal brain regions. Critically, it was largely expression from brain-resident cells in the Adjacent Normal AOIs that significantly increased the hazard of death, more so than the expression from metastatic tumor cells; supporting the previous finding that microglia and astrocytes in adjacent normal brain regions are remodeled to support an immune evasive and fibrotic microenvironment in brain metastases.^21^

Given that little is known about how metastatic cells may co-opt features of the host microenvironment, we set out to explore whether cell types in the BrMet TME could employ shared transcriptional signals relevant to disease progression. With this is in mind, we developed the Equivalent Expression Index as an innovative tool that could identify Equivalently-Expressed Genes (EEGs) across similar and dissimilar AOI types. The Equivalent Expression Index identified EEGs with statistical significance and high specificity by modeling the observed biological data and accounting for accuracy. It should be noted that genes that are equivalently-expressed genes are not synonymous with genes that are *not* differentially expressed. The Equivalent Expression Index provides a framework for identifying shared transcriptional programs across any two groups of interest; serving as a broadly-applicable tool to further understand organotropism and metastatic cell adaptation.

Given that cross-talk between metastatic cells and brain-resident cells may drive brain metastases biology, we focused our identification of EEGs to Malignant and Brain-Enriched AOI types. Using Boolean logic-based set analysis, we identified two novel gene signatures (MIBS-9 and ARCS-81) relevant to the BrMet TME. The MIBS-9 signature – derived from the intersection of genes upregulated by BrMets (Δ_M,P_) and equivalent in the brain ($_A,N_) – represented mechanisms that metastatic breast cancer cells may employ to mimic, or adapt to, the brain microenvironment. MIBS-9 genes were expressed across the normal brain; and each gene has been shown to play a role in brain-specific functions and/or neurodegeneration. For example, *MSMO1* (Methylsterol Monooxygenase 1), which plays a key role in biosynthesis of cholesterol^56^ is highly expressed in fetal and mature astrocytes and may play an important role in brain cholesterol metabolism.^57^ *REEP5* is predominantly expressed in neurons, supporting the role of the REEP family of proteins in regulating cellular stress responses, particularly *REEP1*, which is important for maintaining ER organization in corticospinal neurons.^58^ Additionally, *SYNGR2* (Synaptogyrin 2), is enrichment in endothelial and microglia cells consistent with its proposed role in immune cell infiltration.^59^ As a juxtaposition to the MIBS-9 signature, we derived the ARCS-81 signature to explore genes upregulated by the Adjacent Normal brain (Δ_A,N_) and equivalent in breast cancer ($_P,M_); representing pathways employed by, or activated in, the brain to support the formation and/or presence of the metastatic niche. The ARCS-81 signature suggested brain-specific genes that may mimic, or take on the expression patterns, of the cancer cells found in the brain microenvironment. ARCS-81 genes are highly expressed in the cerebellum, a frequent anatomical sites of BCBM.^60^ This novel gene signature contained brain-associated genes known to play a role in cancer (*NEUROD1*^35,36^, *TSPAN9*^37^*, CAMK2*^40^). Interestingly, only a subset of ARCS-81 genes (e.g., *NONO*, *CTBP2*, *TMX2*) were also expressed in the normal breast, suggesting shared genes exploited by metastatic cells to adapt to the brain microenvironment. For example, *NONO*, which is upregulated in neural stem cells^61^, regulates neuron migration/maturation and is linked to neurodevelopmental disease^62^. In the context of breast cancer, *NONO* promotes the splicing, stabilization, and subsequent expression, of its pro-proliferation targets: *SKP2*, *E2F8*, *Myc*, *Cyclin D1*, *Aurora Kinase A*, enhances *EGFR* transcription, and increases breast cancer cell growth and drug resistance through stabilization of *STAT3*.^63^ Moreover, in the brain CTBP2 is expressed in the subventricular zone where its activation promotes neuronal and glial cell differentiation^64^; while in cancer, CTBP2 mediates the repression of E-cadherin and PTEN, promoting the epithelial-to-mesenchymal transition.^38,39^ Lastly, TMX2 is a key regulator of neuronal proliferation, migration, and organization^65^; and promotes cell proliferation in breast cancer, including upregulating genes related to cell survival and metastasis.^66^ The reported brain– and cancer-specific dual roles of these genes make them intriguing targets to test how brain-resident cells support metastatic breast cancer cells.

When we assessed the association between MIBS-9 and ARCS-81 expression and patient survival, we observed that the signatures exhibited opposing correlations. When metastatic breast cancer cells upregulated brain-associated genes (high MIBS-9 expression), this correlated with improved survival. Conversely, when brain-resident cells upregulated cancer-associated genes (high ARCS-81 expression) this correlated with worsened survival. The ability of metastatic cells to take on the expression profiles and/or phenotypes of host organs has been previously documented. For example, in bone metastases, metastatic breast cancer cells have been shown to express osteoblast-specific markers and bone matrix proteins.^67^ Although mimicry of the host microenvironment may help to explain metastatic seeding, we found that “brain-mimicry” was associated with longer survival. Whether this difference is due to suppression of tumor growth by maintenance of a normal brain phenotype remains to be determined. Although the cause of death in the majority of BrMet patients is due to systemic disease progression^68^, the development of BrMets denotes an inflection point associated with decreased survival. Our data showed adaptation of the brain microenvironment to a tumor-supporting, or even tumor-mimicking microenvironment was correlated with decreased survival. In the context of BCBM, we found the effect of metastatic cells on the adjacent normal brain of critical importance, yet relatively understudied.

We used the Equivalent Expression Index to identify and refine novel, biologically-relevant shared gene expression patterns in the primary breast TME of patients who developed BrMets. Specifically, we identified signatures that captured genes relevant to the brain (BEPS-154) and brain-metastatic potential (BACE-45). Importantly, these two signatures were validated not only with external datasets, but also with BrMet-free and overall survival times, while having 3– and 10.8-times fewer genes than PSS-484 (genes upregulated in Malignant AOIs of Progressor-Primary (P) compared to Non-Progressor-Primary (Q)). With the continued growth of whole transcriptome and spatial –omics data, which can link numerous genes to clinical outcomes, the Equivalent Expression Index offers a way to refine large gene signatures into focused, biologically meaningful targets—without compromising key associations with clinical variables. We outline our stepwise refinement of these gene signatures, highlighting the impact of EEGs in identifying which shared pathways were being retained and likely serving as key drivers of BCBM. We identified 15 gene sets present across all three Progressor-Primary signatures, many of whose genes have been implicated in cancer and metastasis. Intriguingly, this suggests that activation of these pathways in the primary breast TME may facilitate brain metastatic capabilities.

Although our findings provide novel biological insights into BCBM and a potential means of tumor and brain-resident cell adaptation in the TME, we acknowledge several important limitations. Our primary data were acquired from tissue microarray (TMA) cores and may not have captured the heterogeneity of the entire tissue. However, the majority of TMA cores were highly homogenous, as validated both by certified pathologists and by high RNA expression reproducibility across AOIs within the same core (**Supplemental Figure 10C)**. Additionally, we acknowledge that although our AOIs are cell subtype enriched, RNA expression measurements are not on the single cell level. To address this, we validated AOI-derived gene expression data in the only publicly available single-cell RNA sequencing data on BCBM, although this dataset had limited representation of brain-resident cell types (**Supplemental Figure 2B-2F**). Lastly, while we anticipate broad applicability of the Equivalent Expression Index, our current findings are hypothesis-generating and specific to breast-to-brain metastases. Future studies are warranted to validate the novel gene signatures introduced here, including cell and biochemical assays to gain insight into mechanisms of regulation. Additionally, single-cell resolution profiling, particularly in Brain-Enriched AOIs, would be useful to further resolve the brain cell subtypes contributing to gene signature expression. Finally, we recommend expanding our approaches to study brain metastases (and their corresponding primary tumors) from various origins (i.e., lung, melanoma, colorectal) to assess whether these signatures are agnostic to primary tumor type, with the potential to identify unified mechanisms of brain metastatic competence.

In conclusion, our work provides a framework on which to understand the converging roles of brain tropism and microenvironmental support of breast cancer brain metastases. Combining the Equivalent Expression Index with differential expression analysis allowed our exploration of transcriptional programs that may be co-opted by metastatic cells, or acquired by normal brain cells, to facilitate seeding and outgrowth of brain metastases. Our Equivalent Expression Index may serve as an important tool for others to characterize equivalent expression between similar and dissimilar cell types across the TME. Ultimately, this data provide a valuable resource for the exploration and identification of therapeutic targets that may be used to modulate the BrMet TME for patient benefit.

## METHODS

### Human Biospecimen Collection

Human subject aspects of this study were approved by the Stanford Institutional Review Board (IRB) in accordance with the Declaration of Helsinki guidelines for ethical conduct of research. We confirm that written informed consent was obtained from all patients involved. After obtaining IRB approval (IRB-34363, IRB-57236) the Stanford STRIDE database was queried for female patients (>18 years of age) with brain metastases from invasive breast cancer (IBC), seen at Stanford Health Center (SHC) between 2008 and 2019. Patient charts were individually reviewed to confirm radiographic evidence of brain metastasis^69^, as well as breast cancer molecular subtype (based on primary breast tumor hormone receptor (ER/PR) and HER2 overexpression statuses). STRIDE was also used to query for patients with IBC (without brain metastases; confirmed by brain MRI), ductal carcinoma in situ (DCIS), and epilepsy (patients undergoing temporal lobectomy for medically refractory epilepsy). Formalin-fixed paraffin-embedded (FFPE) tissue samples, from surgical resections, for corresponding patients were obtained from Stanford Neuropathology and Pathology. Surgical specimens were assessed and annotated by pathologists with expert experience in breast (R.W.) and brain (H.V.) tissues to confirm disease diagnosis and tissue quality. De-identified FFPE blocks were used to build a tissue microarray (TMA) – TMA593, TMA594, and TMA594 – containing 235 tissue cores (1.5mm) from 149 unique patients (including positive control tissue cores: placenta, spleen, tonsil, normal lymph node). Summary statistics of tissues profiles is available in **Supplemental Table 1**. A representative visualization of AOI-level annotations (TMA593, 95 AOIs) is provided in **Supplemental Figure** Clinical information for patients was obtained via STRIDE and electronic health records. Specifically, breast cancer molecular subtype, dates of diagnosis for primary IBC and brain metastases, dates of surgical resections for primary IBC and brain metastases, presence of metastases outside the central nervous system (“non-CNS” distant metastasis), neoadjuvant chemotherapy, neoadjuvant radiation therapy, survival status, and date of death (or date of last contact if alive or if death could not be confirmed). This information was used in time-to-event analyses and as covariates in regression Cox proportional hazards regression analyses.

### Immunohistochemistry

Sections of 5µm thickness derived from FFPE TMA blocks were cut for histological (H&E) staining and immunohistochemistry (IHC). IHC was performed according to manufacturer instructions (Vector Laboratories) using the Vectastain Elite ABC-HRP Kit (PK-6101). Anti-Pan-Cytokeratin (Novus, NBP2-33200), anti-GFAP (Novus, NBP2-33184), and anti-CD45 (CST, 13917), primary antibodies were used. Representative visualizations of core and AOI level IHC/IF staining is provided in **Supplemental Figure 9A, 9B**. Slides were scanned at 40X resolution.

### Digital Spatial Profiling (DSP)

DSP experiments were conducted according to manufacturer’s instructions: GeoMx-NGS Slide Preparation User Manual (MAN-10115-05), GeoMx-NGS Instrument User Manual (SEV-00087-05 for software v2.4), GeoMx DSP NGS Readout User Manual (MAN-10153-01), and GeoMx-NGS Data Analysis User Manual (SEV-00090-05 for software v2.4), with minor modifications. Briefly 5µm FFPE TMA slides were baked overnight (∼18 hours) at 60°C deparaffinization, rehydration, and antigen retrieval. Slides were incubated for ∼23 minutes at 100°C in Antigen Retrieval Solution using a digital electric food steamer. Slides were incubated at 1µg/m: proteinase K for 15 minutes in at 37°C water bath. To preserve tissue morphology, slides were incubated in 10% neural buffered formalin (NBF) and NBF stop buffer. For in situ RNA probe hybridization, 250µL of hybridization solution (containing 25µL of RNA probe mix) was added to each slide and incubated at 37°C overnight (∼16-18 hours) in a hybridization oven. Slides were stringently washed, then labeled with four morphology marker antibodies: (1) SYTO13-AF488, and (2) anti-Pan-Cytokeratin-AF532 antibodies from the Nanostring solid tumor morphology kit along with (3) anti-GFAP-AF594 (Novus Biologicals, NBP2-33184DL594), and (4) conjugated anti-CD45-AF647 (CST, 13917). Slides were loaded into the DSP instrument and scanned to produce a high-resolution digital image based on immunofluorescent (IF) morphology markers. H&E and IHC staining was used to guide identification of tissue regions. Morphology markers was used to guide selection of Malignant (Pan-Cytokeratin), Brain-Enriched (GFAP), and Immune-Enriched (CD45) areas of illumination (AOIs). Indexing oligos (covalently attached to mRNA hybridization probes) were released from each AOI by exposure to UV light, collected by a micro-capillary tip, and deposited in a 96-well plate as described by [ref (22)]. For each AOI, 18,677 indexing oligos, including 80 negative control oligos were measured.

### Library Preparation and Sequencing

After collection of photocleaved oligos, 96-well plates were sealed with a permeable membrane and dehydrated overnight at room temperature. Samples were rehydrated with nuclease-free water and subjected to sequencing library preparation. Indexing oligos from each AOI, containing a unique molecular identifier (UMI), were PCR amplified us primers (GeoMx Seq Code Primer Plate) that hybridize to constant regions in the oligos as well as contain unique dual-indexed barcodes, preserving AOI identity. After library synthesis, 4µL from each well (per plate) were pooled together and purified using AMPure XP beads (Beckman Coulter, A63880), quality tested via High Sensitivity DNA Bioanalyzer chip (Agilent Technologies) and sequenced on an Illumina NovaSeq 6000 instrument (Novogene). Sequencing was paired-end (2 x 150bp) with an additional 2 x 8bp for index sequences. Raw sequencing reads from FASTQ files were mapped to the Nanostring GeoMx whole transcriptome atlas (Hs_R_NGS_WTA_v1.0) using the GeoMx NGS pipeline software (version 2.3.3.10) and saved as digital count conversion (DCC) files. DCC files, along with the Lab Worksheet annotation file and configuration file, were then used to construct a AOI by gene expression count matrix using the R packages: NanoStringNCTools, GeoMxTools, GeoMxWorkflows.

### DSP Data and AOI Annotations

Sequencing reads were trimmed, stitched, and aligned to the WTA list of indexing oligos to identify the source RNA probe. UMI regions of reads were used for deduplication. Data quality was determined per individual AOI. AOIs were excluded from the dataset based on the following criteria: less than 1000 sequencing reads, less than 80% trimmed, stitched, and aligned sequencing reads, and less than 50% sequencing saturation (**Supplemental Table 2**). A high-confidence gene expression detection threshold was determined using the limit of quantification (LOQ) metric. The LOQ is calculated based on the distribution of negative control probes (raw counts) and is used to approximate the quantifiable limit of gene expression per AOI using the formula:

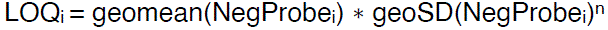

wherein “i” is i^th^ AOI in the dataset and n is the standard deviation value. Per manufacturer’s recommendations, “n” was set to a value of 2, indicating the gene detection threshold is two standard deviations above the geometric mean of the negative probes, for each AOI. A minimum LOQ value of 2 was set as a cutoff. We assayed, on average, 2 AOIs per 1.5mm tissue core (**Supplemental Figure 10A**), and observed high reproducibility, as evidenced by strong correlations (*r* = ≥0.97, p <2.2e-16) in raw counts between replicate AOIs (**Supplemental Figure 10B, 10C, Supplemental Table 4**); indicating the robustness of DSP to reliably detect and quantitate RNA expression. To accurately profile true biological signal, we filtered the gene panel to 9,575 genes which were above the limit of quantitation (LOQ) threshold in at least 1% of AOIs (**Supplemental Figure 10D, Supplemental Table 5**). Genes below the LOQ threshold were excluded from downstream analyses. Each cell type (apart from Stroma-Enriched due to low AOI number), tissue type, and molecular subtype was observed across the range of gene detections rates, indicating that RNA expression profiles are not biased by the sensitivity of DSP gene detection (**Supplemental Figure 10E**). To enable comparisons across AOIs, prior to downstream analyses, raw gene counts were normalized to the upper quartile (Q3 normalization) for each AOI (**Supplemental Figure 11A-D**). Negative probe normalization, background subtraction normalization, and background subtraction followed by Q3 normalization were also tested to ensure the optimal technique for this data (**Supplemental Figure 11D**). In line with previous DSP studies^70^, we observed positive correlations between: AOI surface area and estimated nuclei counts, AOI surface area and raw gene counts, and estimated nuclei counts and raw gene counts (**Supplemental Figure 12A, 12B**). However, after either Q3 normalization or background subtraction followed by Q3 normalization, correlations between surface area/nuclei count and gene counts were significantly reduced. Little to no correlation was observed between AOI surface area/estimated nuclei counts and number of genes detected per AOI. AOI annotations were validated in several ways: (1) Annotations were verified against IF-stained images of each AOI. Labeling AOIs by IF marker abundance aligned with cluster annotations (**Supplemental Figure 2A**). (2) Cell type signatures for Malignant, Brain-Enriched and Immune-Enriched clusters (composed of the 20 most highly expressed genes per cluster) revealed known cell type markers genes (**Supplemental Figure 2B**). (3) Using publicly available single-cell RNA-sequencing data of brain metastases^23^, subset to primary breast cancer (**Supplemental Figure 2C**), we found that Malignant, Brain-Enriched and Immune-Enriched AOI-derived cell-type signatures were highly enriched in metastatic-tumor, astrocyte, and immune (macrophage/T cell/dendritic) cell populations; respectively (**Supplemental Figure 2D**), corroborating our cell type annotations. (4) Lastly, anticipating that Immune– and Brain-Enriched AOIs likely represent an admixture of cells, we used spatial deconvolution to estimate the proportion of immune and brain cell types, respectively, in these AOIs (**Supplemental Figure 2E, 2F**). Immune-Enriched AOIs contained high proportions of stromal, myeloid, and T/NK cells; while Brain-Enriched AOIs were largely composed of neurons and pericytes.

### Differential Expression (DE) Analysis

Filtered and normalized AOI-level count data was used for DE analyses to identify specific RNA differences between tissue types and cell types (Supplemental File 2). Per Nanostring recommendations, to account for tissue subsampling, a linear mixed effects model was used for analysis, with TMA core ID as the random effect (GeoMx Tools R Package). Genes with abs(Log_2_FoldChange) ≥ 0.5 along with a p-value ≤ 0.05 were considered differentially expressed.

### Gene Set Enrichment Analysis (GSEA)

GSEA was carried out as previously described.^71^ Gene sets used in this study include Human MSigBD hallmark, Human MSigBD gene ontology (biological process, cellular component, molecular function), and 70 gene sets curated for this study (**Supplemental File 2**). GSEA results available in **Supplemental File 3**.

### Equivalent Expression Index

We take an analytic approach to identify genes with “equivalent” expression between groups. The idea for the Equivalent Expression Index is based on an Empirical Bayes method.

Specifically, suppose that the between group difference for gene’ can be estimated by 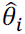, which follows a normal distribution, whose standard deviation can be estimated by *σ_i_*,

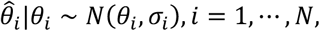

where *θ_i_* is the true between-group difference, which is unknown to us. The objective of the Equivalent Expression Index is to identify genes such that |*θ_i_*| ≤ *δ*_0_ for a given equivalence threshold *δ*_0_. In this study, the gene expression level is Log_2_ transformed, and we set *δ*_0_ = 0.2, thus, the “equivalence” gene set is defined as genes met the following criterion:

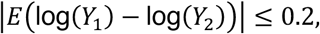

where *Y_j_* is the expression level under condition *j* ∈ {0,1}. Using the approximation,

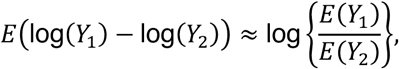

this equivalence criterion is approximately the same as

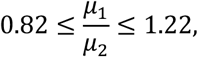

where *µ_j_* is the mean expression level under condition *j* ∈ {0,1}. In summary, our objective is to identify genesfrom the set:

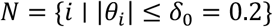

To this end, we assume that:

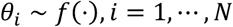

Based on 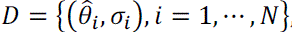, we can estimate the density function of *f*(⋅). Denote the resulting estimator by 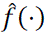, the posterior distribution of *θ_i_* s:

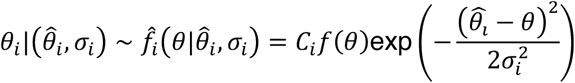

where *c_i_* is a normalizing constant. We can then construct a credible interval for *θ_i_* based on 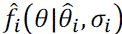 such that:

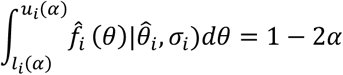

where α is a given threshold to ensure that 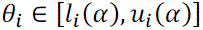 with a probability no smaller than 1 − 2α. The equivalence gene can be identified as:

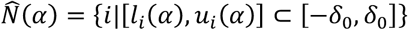

The “accuracy” of identified equivalence genes can be measured by:

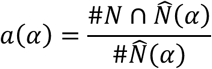

We want to select α to ensure that this “accuracy” is high enough, i.e., α(α) > 95%. In practice, we need to estimate this precision empirically from the data. To this end, we use Monte-Carlo method. Specifically, we can

(1) Simulate “true” between-group difference 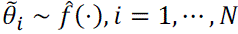,
(2) Simulate observed between-group difference based on the “true” differences 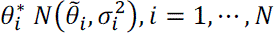,
(3) Using the proposed method to construct 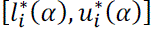 based on simulated data 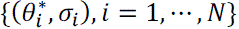
(4) Calculate 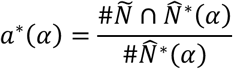
(5) where 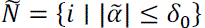 and 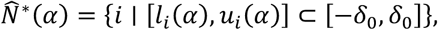
(6) which is equivalence gene set estimated from the simulated data.

This accuracy-adjusted approach to identify EEGs, takes into account the observed data (based on estimated between group differences 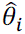) to generate an estimated distribution model for *θ_i_* For example, when EEGs are identified between two similar AOI types (e.g., Malignant AOIs from Progressor-Primary (**P**) and Malignant AOIs from Metastasis (**M**) the estimated *θ_i_* distribution has an increased proportion of genes within the (–0.2, 0.2) interval; whereas analysis between dissimilar AOI types (e.g., Malignant AOIs from Progressor-Primary (**P**) and Brain-Enriched AOIs from Normal Brain (**N**)) have a lower proportion of genes in this interval (**Supplemental Figure 4A, 4B**). To validate the ability of the Equivalent Expression Index to accurately identify EEGs, we performed t-testing on five randomly selected genes from each pairwise comparison, a total of 30 genes. We set the accuracy threshold to 5% FDR, such that we expect of the 30 genes identified, 2 genes may be “false positive” EEGs, and 28 should be “true positive” EEGs (**Supplemental Figure 4C**). As expected, we find that the normalized expression of the 30 EEGs identified show no significant difference between groups, except for two genes: *DVL2*, *MAPK1* – confirming the accuracy of our approach (**Supplemental Figure 4D**). Additionally, we used bootstrapping (500 iterations) to statistically test each EEG set across all pairwise comparisons (**Supplemental Figure 4E**). Importantly, the EEG set corresponding to the specific pairwise comparison that produced them showed the lowest proportion of significant p-values, indicating the highest proportion of insignificant p-values. This outcome, influenced by the presence of EEGs, underscores the specificity of this novel approach. Equivalent Expression Index data available in **Supplemental File 4**. Gene signatures derived in this study are available in **Supplemental File 5**.

### Overall Expression Enrichment

Gene signature enrichment scores were calculated using the “Gene Set Overall Expression (OE)” method as described by Yeh et al.^72^ This method computes gene signature single sample scores. Briefly, normalized DSP count data was transformed to transcripts per million. Then, OE scores for a given signature, based on a user-inputted gene list, was computed for each AOI. OE scores served as a readout of the relative enrichment of a signature given the expression levels of associated genes in the AOI.

### Spatial Omics Overlay

Gene signature scores were overlaid onto AOIs using the SpatialOmicsOverlay R package (Nanostring). Briefly, signature scores for each AOI were computed as described above. The Lab Worksheet annotation file and OME-TIFFs (for each DSP scan) were loaded into R and AOIs were overlaid onto the high-resolution digital image using the “readSpatialOverlay” function. AOIs were then visualized using metadata factors (i.e., cell type, signature scores, etc.).

### Statistical Analysis

Statistical test for data analysis included Welch two-sample t-test, two-sample Wilcoxon test, ANOVA followed by Tukey HSD test, log-rank test, and Student’s t-test. For Gene Set Enrichment Analysis (GSEA), genes were ranked based on Log_2_FoldChange from differential expression analysis. Pre-ranked GSEA was conducted using the GSEA software (v4.3.2).^71^ Genes were mapped to the Hallmark gene sets, gene ontology biological processes, cellular components, and molecular functions pathways obtained from the Molecular Signatures Database (MSigDB) Human Collection (v2023.2). Ranked genes were also mapped to a curated dataset (n = 70 gene lists) enriched for breast– and brain-related gene sets. Univariate and multivariate cox proportional hazards regression was used to assess the independent association of gene signatures scores with time-to-event as well as their 95% confidence intervals. Data for cox proportional hazards models consisted of Overall Expression gene signature scores and patient clinical information (death status, brain metastases status, days from brain metastases resection to death (or last contact), days from primary breast cancer diagnosis to brain metastases diagnosis, and days from primary breast cancer diagnosis to death (or last contact)). Forest plots and associated regression statistics were generated using the ‘survival’ and ‘survminer’ packages in R. In all cases, differences were considered statistically significant when p-value was less than 0.05. In all cases, differences were considered statistically significant when p-value was less than 0.05. All graphical representation and statistical analyses of data were performed using R. Images panels were assembled in Adobe Illustrator with brightness and contrast modifications made only for image clarity.

### Data Availability

All data collected in this study, including GeoMx digital spatial profiling and deidentified clinical metadata will be deposited and made publicly accessible through Zenodo. Upon publication raw sequencing files and high-resolution slide scan files (OME.tiff) will be available via Synapse, due to the large number of files and their substantial size. This deposition will also include raw and Q3 normalized count matrices, LabWorksheet.csv, Probe Kit Configuration (PKC) file, Biospecimen metadata file, and DCC count files. Expression data (Raw and Q3 normalized count tables) and AOI-level metadata (Sample ID, Patient ID, Tissue Type, Cell Type, and Molecular Subtype) are available in Supplemental File 1. The single cell RNA sequencing brain metastases data^23^ used in this study were accessed from GEO (GSE186344). The microarray expression studies with breast tumor samples were accessed from the GEO (GSE194040, GSE2034, and GSE102484). Further information and requests for reagents and resources should be directed to, and will be fulfilled by, the lead contact, M.H.G. This study did not generate new unique reagents or cell lines.

### Code Availability

All data processing and analysis code will be available via GitHub once paper is published. The R script used in this study for the Equivalent Expression Index will also be deposited and made publicly accessible through Zenodo. The GitHub repository includes documented code to perform the Equivalent Expression Index analysis including README.md, license, Equivalent_Expression_Index.Rmd, and example input and output datasets (Input and Output folders) for testing the R script, aligned with the steps outlined in README.md.

## ACKNOWLEDGMENTS

The authors thank M.A. for use of the GeoMx DSP instrument. The authors thank Nanostring for technical support and for providing R vignettes for specific portions of data analysis. The authors thank Stanford Health Center patients (and their families) for donating tissue for research purposes.

## AUTHOR CONTRIBUTIONS

M.UG., S.K.P., and M.HG. designed the study. M.UG. performed the spatial profiling experiments with the supervision of M.A. and M.HG. B.G. identified patient tissue samples for study. S.A., A.T., and H.V. collected and annotated archival brain metastases and normal brain tissue samples. R.W. and S.V. collected and annotated archival DCIS, primary breast cancer, and control tissue samples. S.V. constructed tissue microarrays. M.UG., B.G., P.NP., and M.G. collected patient clinical data. M.UG. and C.Y.Y. performed the computational and statistical analyses. S.K.P. and L.T. conceptualized the Equivalent Expression Index. L.T. wrote the Equivalent Expression Index code with input from M.UG. M.UG., S.K.P., and M.HG. interpreted the results. G.B. created data portal and supported computational and statistical analyses. T.T. provided additional support on data collection. M.UG., S.K.P., and M.HG. wrote the manuscript. S.K.P. and M.HG. supervised the study. All authors reviewed and approved the paper.

## SUPPLEMENTAL DATA – TABLES, FIGURES, & FILES

### SUPPLEMENTAL TABLES

**Supplemental Table 1.** Information on site and molecular subtypes of tissues, including control tissues, used for GeoMx DSP study. AOIs were selected from across three Tissue Microarray (TMA) slides: TMA593 (n = 26 cores), TMA594 (n = 104 cores), TMA595 (n = 104 cores). DCIS: Ductal carcinoma in situ.

| Tissue Microarray Cohort (n = 235 cores) |  |  |  |  |  |
| --- | --- | --- | --- | --- | --- |
| TMA593 (n = 26), TMA594 (n = 104), TMA595 (n = 104) |  |  |  |  |  |
|  | Lum A | Lum B | HER2+ | TNBC | --- |
| **Brain Metastasis** | 17 | 23 | 18 | 21 | --- |
| **Adj. Normal**<br>(Brain Metastasis) | 10 | 13 | 10 | 15 | --- |
| **Primary Breast**<br>(Invasive Breast Cancer) | 5 | 3 | 2 | 36 | --- |
| **DCIS** | 1 | 1 | --- | 13 | --- |
| **Local Recurrence /**<br>**Lymph Node** | 1 | --- | --- | 5 | --- |
| **Normal Brain** |  |  |  |  | 23 |
|  | Placenta | Spleen | Tonsil | Lymph Node |  |
| **Controls** | 3 | 5 | 5 | 5 |  |

**Supplemental Table 2.** Sequencing quality control (QC) metrics for AOIs. Table shows the number of AOIs that passed or produced warnings for each QC parameter. QC parameter thresholds were based on Nanostring guidelines. All flagged AOIs were removed from downstream analysis.

| Sequencing Quality Control Metrics |  |  |  |
| --- | --- | --- | --- |
| AOI (n = 473) |  |  |  |
|  | PASS | WARNING | % |
| **Minimum # of Reads (> 1000)** | 472 | 1 | 0.2 |
| **% Trimmed Reads (80%)** | 471 | 2 | 0.4 |
| **% Stitched Reads (80%)** | 457 | 16 | 3.4 |
| **% Aligned Reads (80%)** | 454 | 19 | 4.0 |
| **% Sequencing Saturation (50%)** | 469 | 4 | 0.8 |
| **Minimum Area (> 1000um<sup>2</sup>)** | 473 | 0 | 0 |
| **Maximum NTC Counts (< 1000)** | 473 | 0 | 0 |
| **Minimum Negative Probe (n > 1)** | 473 | 0 | 0 |
| **Total Flagged AOIs**<br>(removed from analysis) | --- | 23 | 4.9 |

**Supplemental Table 3.** Equivalently-Expressed Genes (EEGs) from all pairwise comparisons between AOI types analyzed. Table includes the number of EEGs, percentage of post-QC filtered gene set (n = 9575). Met: Metastasis. AdjN: Adjacent Normal.

| Equivalence Expression Index |  |  |  |  |  |
| --- | --- | --- | --- | --- | --- |
| Comparison |  | Stringent Approach |  | FDR-Guided Approach |  |
|  |  | No. of EEGs | % FGS* | No. of EEGs | % FGS* |
| **EA #1** | Progressor (P) vs. Brain Met (M) | 171 | 1.79 | 3590 | 37.5 |
| **EA #2** | AdjN (A) vs. Normal Brain (N) | 315 | 3.29 | 4095 | 42.8 |
| **EA #3** | Brain Met (M) vs. AdjN (A) | 1095 | 11.4 | 2333 | 24.4 |
| **EA #4** | Brain Met (M) vs. Normal Brain (N) | 157 | 1.64 | 531 | 5.55 |
| **EA #5** | Progressor (P) vs. AdjN (A) | 38 | 0.39 | 70 | 0.73 |
| **EA #6** | Progressor (P) vs. Normal Brain (N) | 8 | 0.08 | 65 | 0.68 |

\*FGS: Filtered Gene Set (9575 genes after filtering by % detection)

**Supplemental Table 4.** Table lists 13 pairs of replicate AOIs (from within the same tissue core) and the Pearson correlation coefficient (based on raw counts) between each replicate AOI.

| Gene Expression Correlation between Replicate AOIs |  |  |  |  |  |
| --- | --- | --- | --- | --- | --- |
| AOIs (n = 26, 13 pairs) |  |  |  |  |  |
|  | Tissue Type | Subtype | Cell Type | r | p-value |
| AOI 37 -- AOI 38 | Metastasis | Lum A | Malignant | 0.9936004 | < 2.2e<sup>-16</sup> |
| AOI 43 -- AOI 44 | Progressor | Lum A | Malignant | 0.9946146 | < 2.2e<sup>-16</sup> |
| AOI 11 -- AOI 56 | Metastasis | Lum B | Malignant | 0.9865149 | < 2.2e<sup>-16</sup> |
| AOI 65 -- AOI 66 | Metastasis | Lum B | Malignant | 0.9851823 | < 2.2e<sup>-16</sup> |
| AOI 26 -- AOI 92 | Metastasis | HER2+ | Malignant | 0.9558926 | < 2.2e<sup>-16</sup> |
| AOI 35 -- AOI 36 | Adjacent Normal | HER2+ | Brain-Enriched | 0.9484561 | < 2.2e<sup>-16</sup> |
| AOI 15 -- AOI 18 | Progressor | TNBC | Malignant | 0.9855439 | < 2.2e<sup>-16</sup> |
| AOI 67 -- AOI 68 | Normal Brain | n/a | Brain-Enriched | 0.9921995 | < 2.2e<sup>-16</sup> |
| AOI 69 -- AOI 70 | Normal Brain | n/a | Brain-Enriched | 0.9601988 | < 2.2e<sup>-16</sup> |
| AOI 72 -- AOI 73 | Normal Brain | n/a | Brain-Enriched | 0.9880215 | < 2.2e<sup>-16</sup> |
| AOI 71 -- AOI 74 | Normal Brain | n/a | Brain-Enriched | 0.9857232 | < 2.2e<sup>-16</sup> |
| AOI 79 -- AOI 80 | Spleen | n/a | Immune-Enriched | 0.9897301 | < 2.2e<sup>-16</sup> |
| AOI 81 -- AOI 82 | Lymph Node | n/a | Immune-Enriched | 0.9732985 | < 2.2e<sup>-16</sup> |

**Supplemental Table 5.** Percent detection of DSP across AOIs. Table shows the percentage of genes detected in a specific percentage of AOIs. Limit of detection (LOD) threshold is set to 2 per Nanostring recommendations.

| % Gene Detection |  |  |
| --- | --- | --- |
| AOIs (n = 450) |  |  |
| % of AOIs | LOD* = 2 |  |
|  | No. of Genes | % of WTGS** |
| 1% (4) | 9575 | 51.3 |
| 5% (22) | 5781 | 31.0 |
| 10% (45) | 4143 | 22.2 |
| 20% (90) | 2607 | 14.0 |
| 30% (135) | 1814 | 9.7 |
| 50% (225) | 893 | 4.8 |
| 75% (337) | 282 | 1.5 |
| 100% (450) | 6 | 0.03 |

\*LOD: Limit of Detection
\*\*WTGS: Whole Transcriptome Gene Set (18677 genes in probe mix)

### SUPPLEMENTAL FIGURES

**Supplemental Figure 1.**
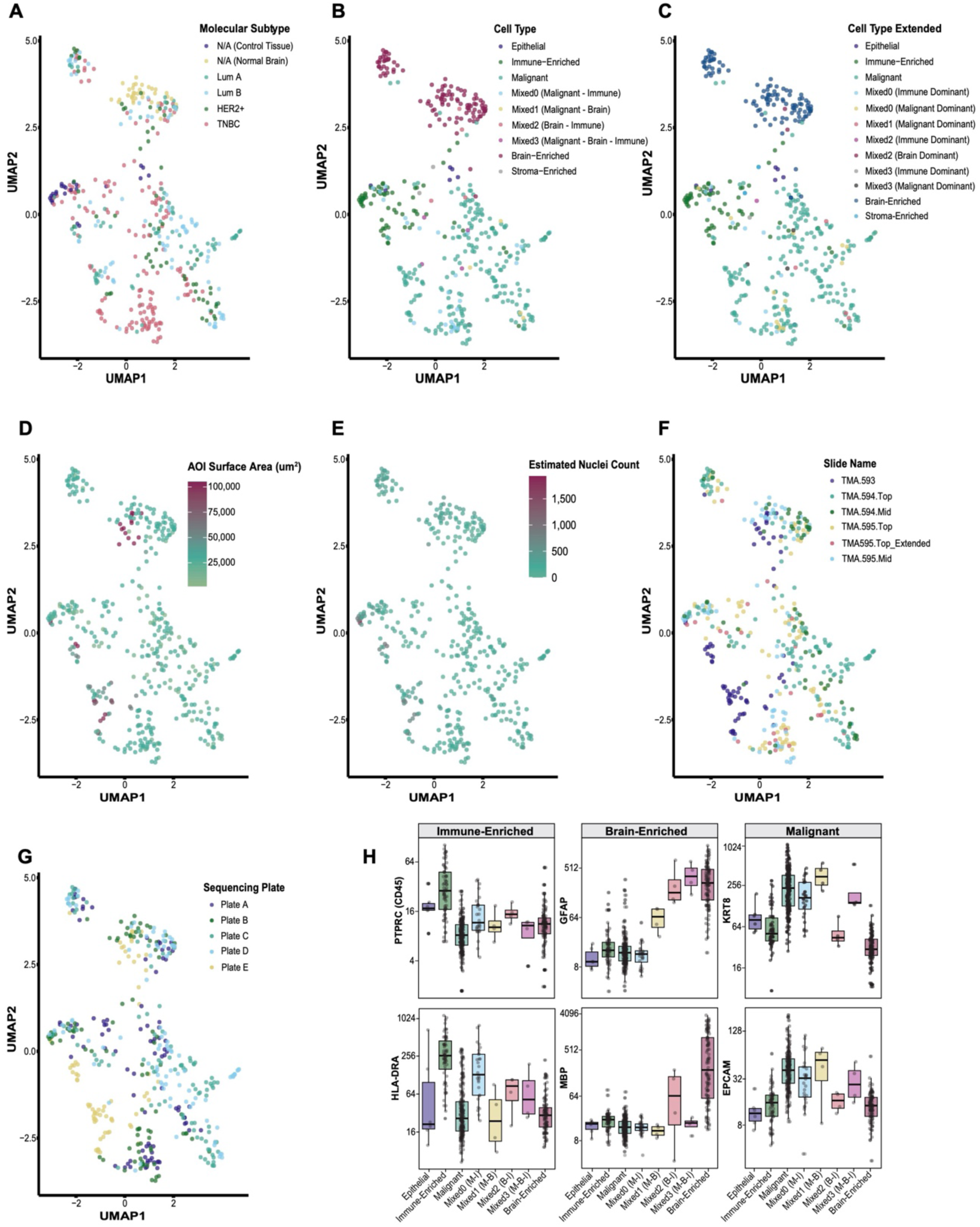
(Supportive Data to Main Figure 2). AOI clustering and cell type-specific gene expression. **(A-G)** UMAP clustering of AOIs annotated by **(A)** breast cancer molecular subtype, **(B)** cell type, **(C)** extended cell type, **(D)** AOI surface area, **(E)** estimated nuclei count, **(F)** slide, and **(G)** sequencing plate. **(H)** Boxplots of Q3 normalized gene counts by AOI extended cell type for selected immune-, brain-, and tumor-associated genes. n = 6, 69, 224, 28, 4, 4, 4, and 109 AOIs in Epithelial, Immune-Enriched, Malignant, Mixed0 [Malignant-Immune], Mixed1 [Malignant-Brain], Mixed2 [Brain-Immune], Mixed3 [Malignant-Brain-Immune], and Brain-Enriched, respectively. For boxplots in **H**, middle line denotes the median, box edges indicate the 25th and 75th percentiles, and whiskers extend to the most extreme points that do not exceed ±1.5 times the interquartile range (IQR). P values are based on nonparametric test (Kruskal–Wallis) followed by Dunn test for pairwise comparisons.

**Supplemental Figure 2.**
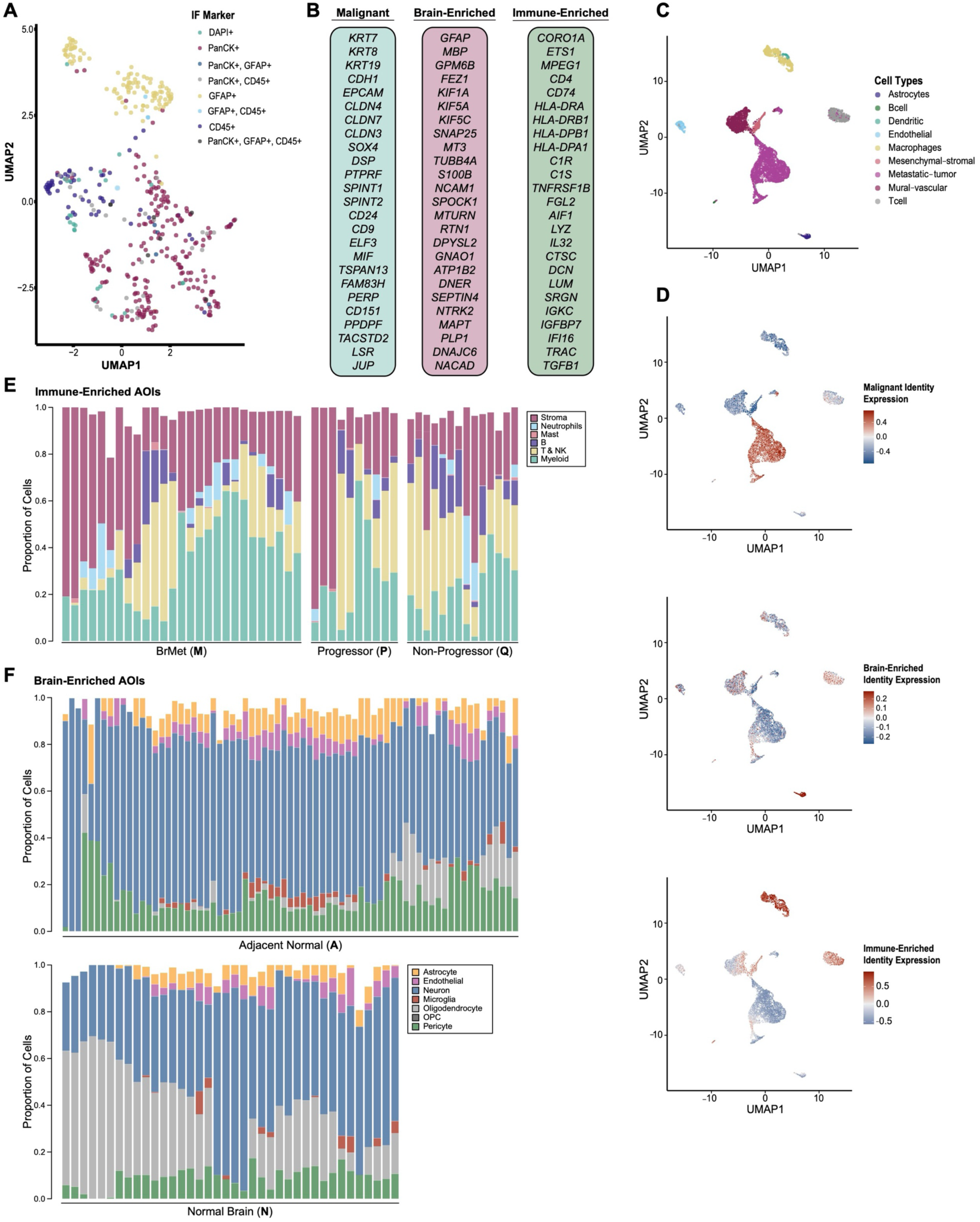
(Supportive Data to Main Figure 2). Validation of AOI annotations. **(A)** UMAP clustering of AOIs annotated by immunofluorescence (IF) marker. **(B)** List of top 25 genes in Malignant, Brain-Enriched, and Immune-Enriched AOIs. **(C-D)** UMAP clustering of single cell RNA-sequencing data of breast cancer brain metastases samples (n = 3, ref. [23]) annotated by **(C)** cell types and **(D)** Malignant, Brain-Enriched, and Immune-Enriched identity signature enrichment expression based on top 25 genes in **B**. **(E-F)** Stacked barplots showing the cell type composition (cell proportion) in each AOI for **(E)** Immune-Enriched and **(F)** Brain-Enriched AOIs.

**Supplemental Figure 3.**
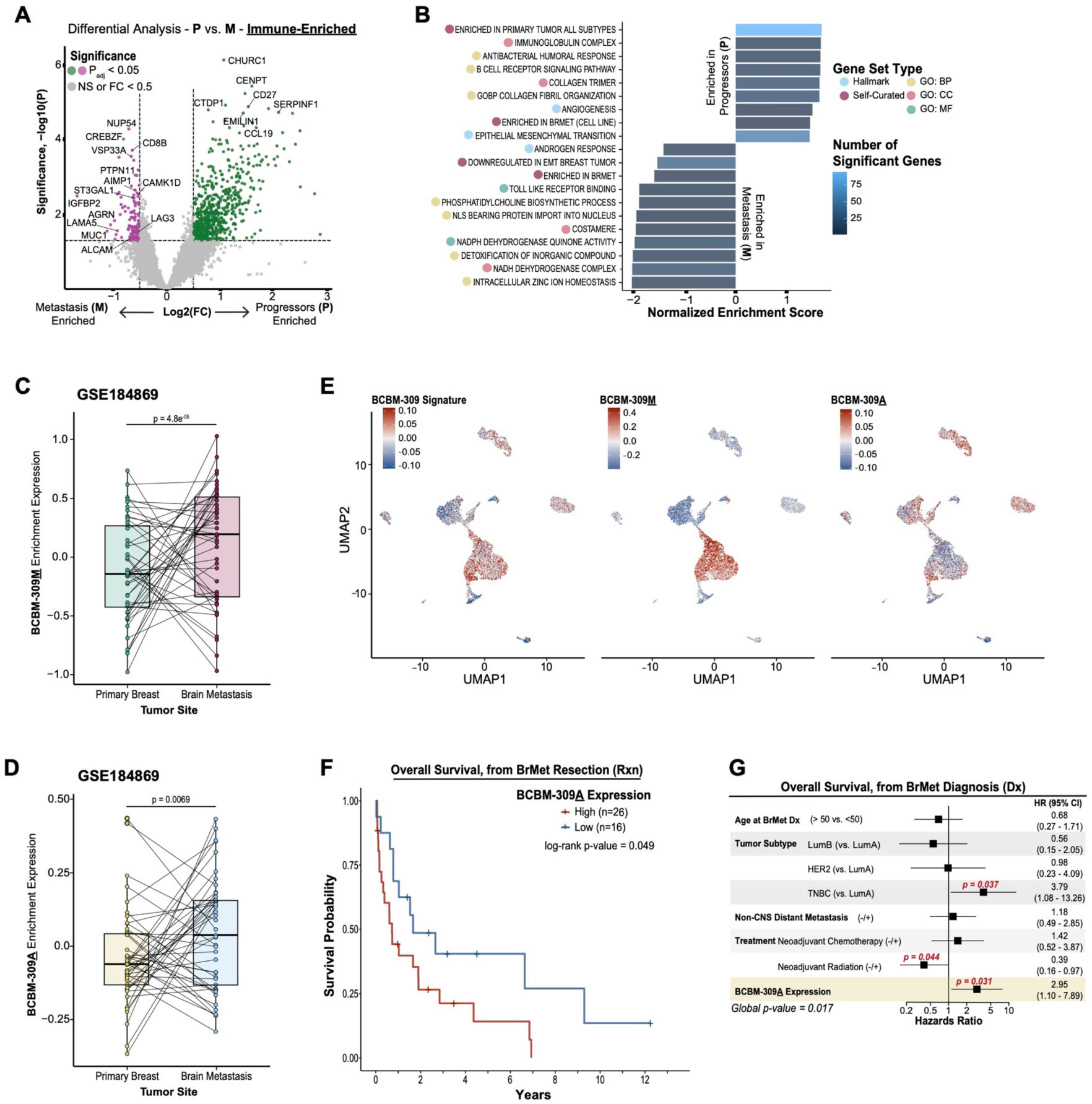
(Supportive Data to Main Figure 3). Differential expression analysis in Progressor-Primary (P) vs Metastasis (M) Immune-Enriched AOIs and BCBM-309 signature in the BrMet microenvironment. **(A)** Volcano plot of differentially expressed genes between Progressor-Primary (n = 10) and Metastasis (n =28) Immune-Enriched AOIs. P-values were calculated using a mixed-effects model using chi-square (χ2) tests (two-sided) with p < 0.05 considered as significant. Genes with p > 0.05 and/or Log_2_(Fold Change) < 0.5 are considered not significant. **(B)** Gene set enrichment analysis of genes upregulated in Progressor-Primary compared to Metastasis Immune-Enriched AOIs. Gene set type and number of significant (p < 0.05) genes are denoted. **(C-D)** Boxplot of **(C)** BCBM-309M and **(D)** BCBM-309A enrichment expression by tumor site. (*Left*) n = 45 and 45 samples from matched primary breast and brain metastasis; respectively (data from ref. [21]). (*Right*) n = 16 and 16 samples from matched primary breast and brain metastasis; respectively (data from ref. [22]). Light grey lines connect paired samples. For boxplots in **C** and **D**, middle line denotes the median, box edges indicate the 25th and 75th percentiles, and whiskers extend to the most extreme points that do not exceed ±1.5 times the interquartile range (IQR). P values are based on paired Wilcoxon rank sum test. **(E)** UMAP clustering of single cell RNA-sequencing data of breast cancer brain metastases samples (n = 3, ref [23]) annotated by BCBM-309, BCBM-309M, and BCBM-309A signature enrichment expression. **(F)** Kaplan-Meier curves of overall survival (from date of brain metastases resection) as a function of BCBM-309A expression. Patients were segregated into two groups (High, n = 26; Low, n = 16) based on the enrichment expression of the BCBM-309A signature in Adjacent Normal AOIs. The log-rank p-value was derived from comparing discretized predictors (high versus low BCBM-309A expression). **(G)** Multivariable Cox proportional hazards regression analysis assessing the impact of clinical covariates and the expression of BCBM-309A on overall survival (from date of brain metastasis diagnosis). In **G**, for each covariate, the hazard ratio (HR) and its 95% confidence interval (CI), denoted by bars, are presented. In **G**, p-values for each covariate were calculated using the Wald statistic. p < 0.05 are significant and denoted in bold red text. p-values near significance are also noted. Global p-value was calculated using the log-rank statistic. FC: fold change. GO: Gene ontology. BP: biological process. CC: cellular component. MF: molecular function. Rxn: Resection. Dx: Diagnosis.

**Supplemental Figure 4.**
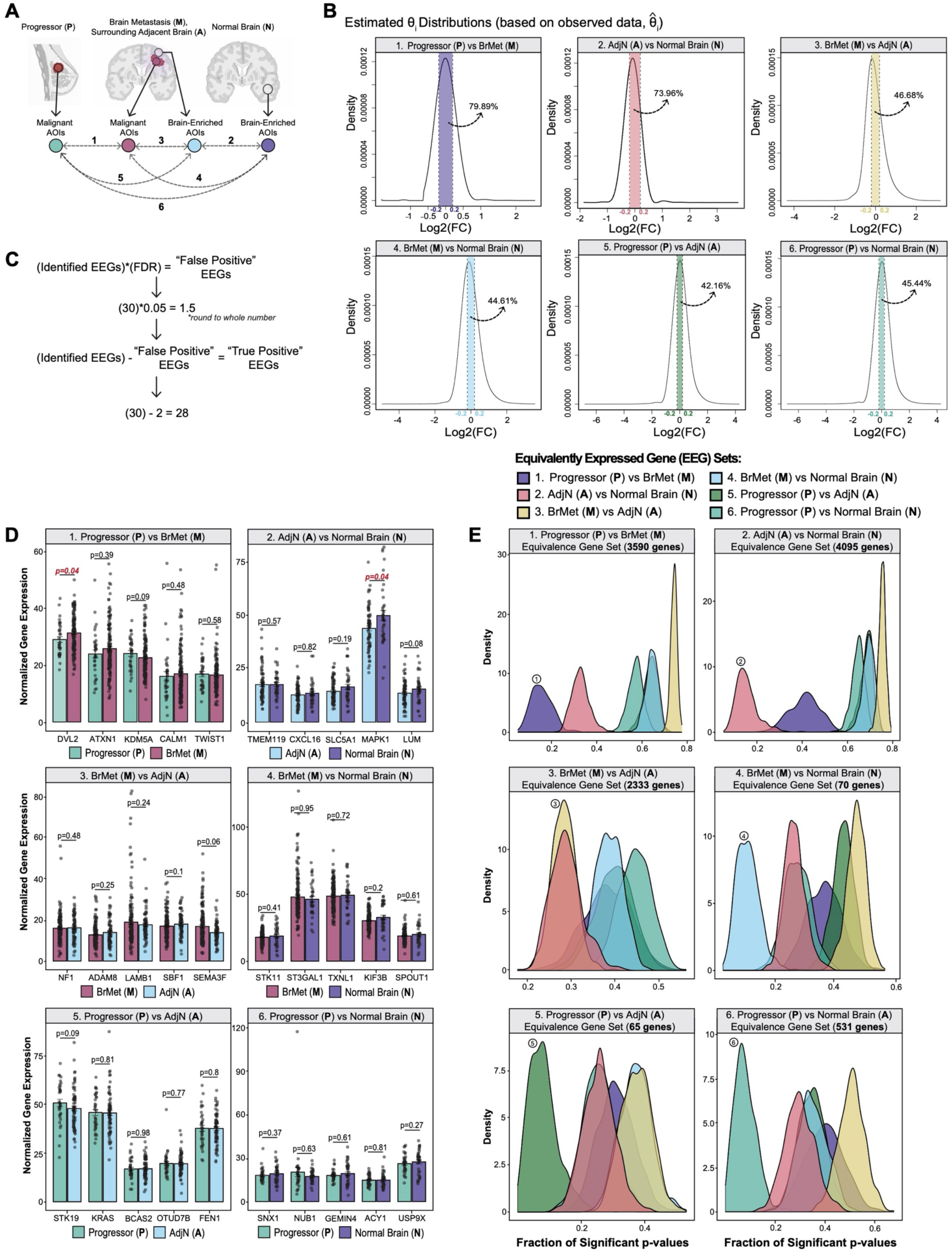
(Supportive Data to Main Figure 4). Validation of the Equivalent Expression Index. **(A)** Schematic of the six pairwise equivalent expression analyses carried out between AOI types: (1) Malignant AOIs from Progressor-Primary (P) versus Malignant AOIs from Metastasis (M), (2) Brain-Enriched AOIs from Adjacent Normal (A) versus Brain-Enriched AOIs from Normal Brain (N), (3) Malignant AOIs from Metastasis (M) versus Brain-Enriched AOIs from Adjacent Normal (A), (4) Malignant AOIs from Metastasis (M) versus Normal Brain (N), (5) Malignant AOIs from Progressor-Primary (P) versus Brain-Enriched AOIs from Adjacent Normal (A), (6) Malignant AOIs from Progressor-Primary (P) versus Brain-Enriched AOIs from Normal Brain (N). Created with BioRender.com. **(B)** Density plots of the estimated *θ_i_* distribution (based on observed 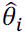 data) for the six pairwise comparisons outline in **A**. X-axis shows the *θ_i_* (equal to the Log_2_(FC) between AOI types). Y-axis shows the density of genes. The area under the curve, representing genes whose Log_2_(FC) between AOI types being compared falls within the Equivalence threshold (δ_0_) [ (–0.2, 0.2)], is highlighted for each plot. The corresponding percentage of genes within this interval is reported within each plot. **(C)** Example schematic for calculating true positive EEGs when FDR is set to 5% (0.05). Calculations correspond to analysis represented in **D**. **(D)** Barplots depicting normalized gene expression of randomly selected genes identified as EEGs by the Equivalent Expression Index for the six pairwise comparisons outline in **A** (n = 30 total EEGs, 5 per comparison group). For barplots in **D**, bar denotes mean expression with ± standard error of the mean (S.E.M.). The p-value of each gene comparison is based on paired Wilcoxon rank sum test. **(E)** Density plots depicting the distribution of significant (p < 0.05) p-values from statistically testing each EEG set across the six pairwise comparisons outline in **A**. For each plot, the p-value distributions from all six pairwise comparisons are depicted. The density plot corresponding to the EEG set of interest (#1-6) is labeled. EEGs: Equivalently Expressed Genes. FC: fold change. FDR: false discovery rate.

**Supplemental Figure 5.**
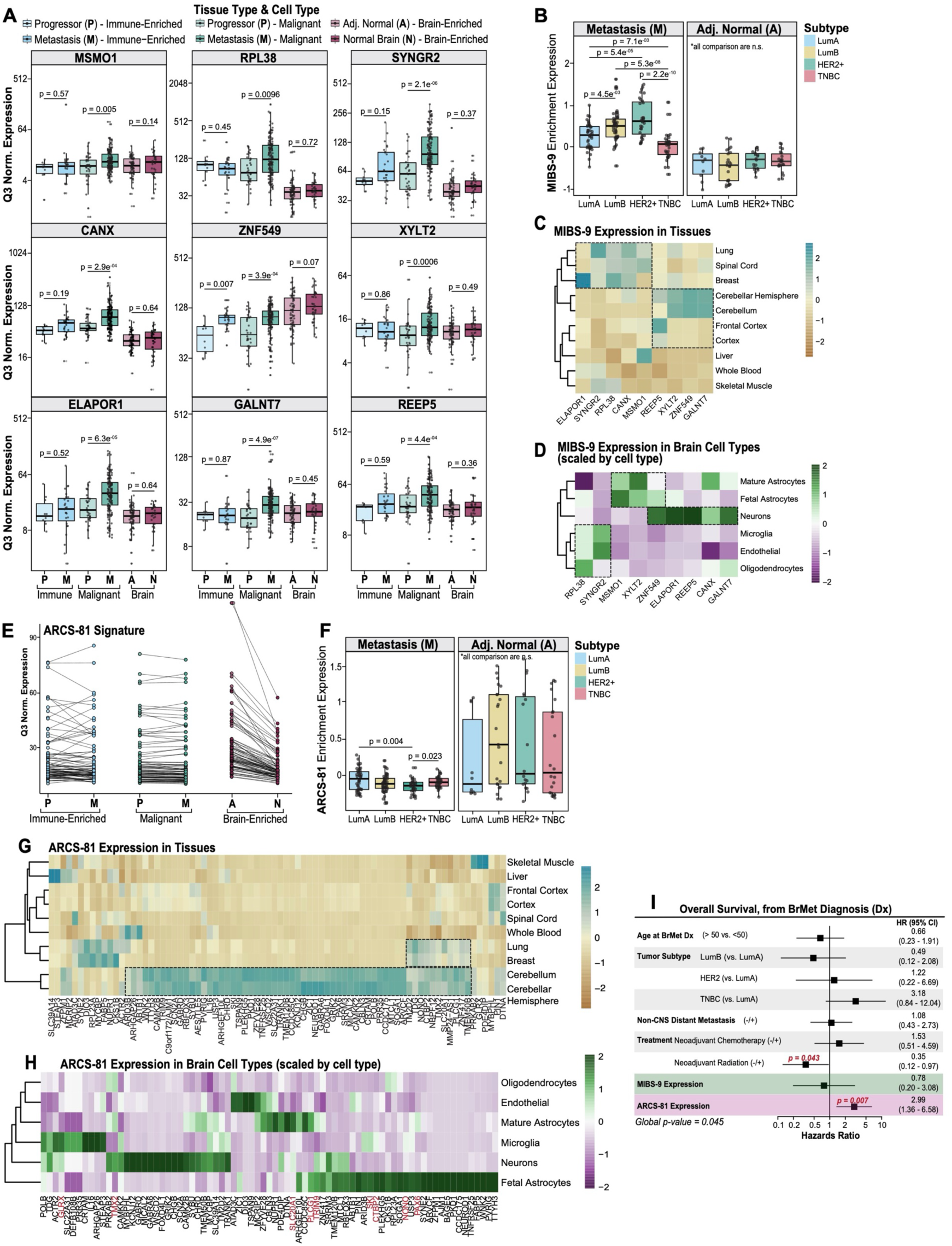
(Supportive Data to Main Figure 5). MIBS-9 and ARCS-81 signatures in the BrMet microenvironment. **(A)** Boxplots of Q3 normalized gene counts of individual genes from the MIBS-9 signature (*MSMO1*, *RPL38*, *SYNGR2*, *CANX*, *ZNF549*, *XYLT2*, *ELAPOR1*, *GALNT7*, *REEP5*), by tissue type and cell type. n = 10, 27, 34, 154, 73, and 38 AOIs in Progressor/Immune-Enriched, Metastasis/Immune-Enriched, Progressor/Malignant, Metastasis/Malignant, Adjacent Normal/Brain-Enriched, Normal Brain/Brain-Enriched; respectively. **(B)** Boxplots of MIBS-9 enrichment expression by breast cancer molecular subtype. n = 154 and 73 AOIs in Metastasis (Malignant) and Adjacent Normal (Brain-Enriched); respectively. **(C)** Heatmap of MIBS-9 signature genes in human tissues (data from GTEx (ref. [33])). **(D)** Heatmap of MIBS-9 signature genes in human brain cell types (data from BrainRNAseq.org (ref. [34])). **(E)** Dot plot of Q3 normalized gene counts of individual genes from the ARCS-81 signature by tissue type and cell type. n = 10, 27, 34, 154, 73, and 38 AOIs in Progressor/Immune-Enriched, Metastasis/Immune-Enriched, Progressor/Malignant, Metastasis/Malignant, Adjacent Normal/Brain-Enriched, Normal Brain/Brain-Enriched; respectively. Each dot represents the normalized expression of one gene. **(F)** Boxplots of ARCS-81 enrichment expression by breast cancer molecular subtype. n = 154 and 73 AOIs in Metastasis (Malignant) and Adjacent Normal (Brain-Enriched); respectively. **(G)** Heatmap of ARCS-81 signature genes in human tissues (data from GTEx (ref. [33])). **(H)** Heatmap of ARCS-81 signature genes in human brain cell types (data from BrainRNAseq.org (ref. [34])). **(I)** Multivariable Cox proportional hazards regression analysis assessing the impact of clinical covariates and the expression of MIBS-9 and ARCS-81 on overall survival (from date of brain metastasis diagnosis). For each covariate, the hazard ratio (HR) and its 95% confidence interval (CI), denoted by bars, are presented. The p-values for each covariate were calculated using the Wald statistic. p < 0.05 are significant and denoted in bold red text. p-values near significance are also noted. Global p-values were calculated using the log-rank statistic. For boxplots in **A**, **B**, and **F**, middle line denotes the median, box edges indicate the 25th and 75th percentiles, and whiskers extend to the most extreme points that do not exceed ±1.5 times the interquartile range (IQR). P values are based on nonparametric test (Kruskal–Wallis) followed by Dunn test for pairwise comparisons. In **C** and **G**, heatmaps are colored from gold to teal according to average expression value of –2 to 2. In **D** and **H**, heatmaps scaled by row, and colored from purple to green according to average expression value of –2 to 2. Dx: diagnosis.

**Supplemental Figure 6.**
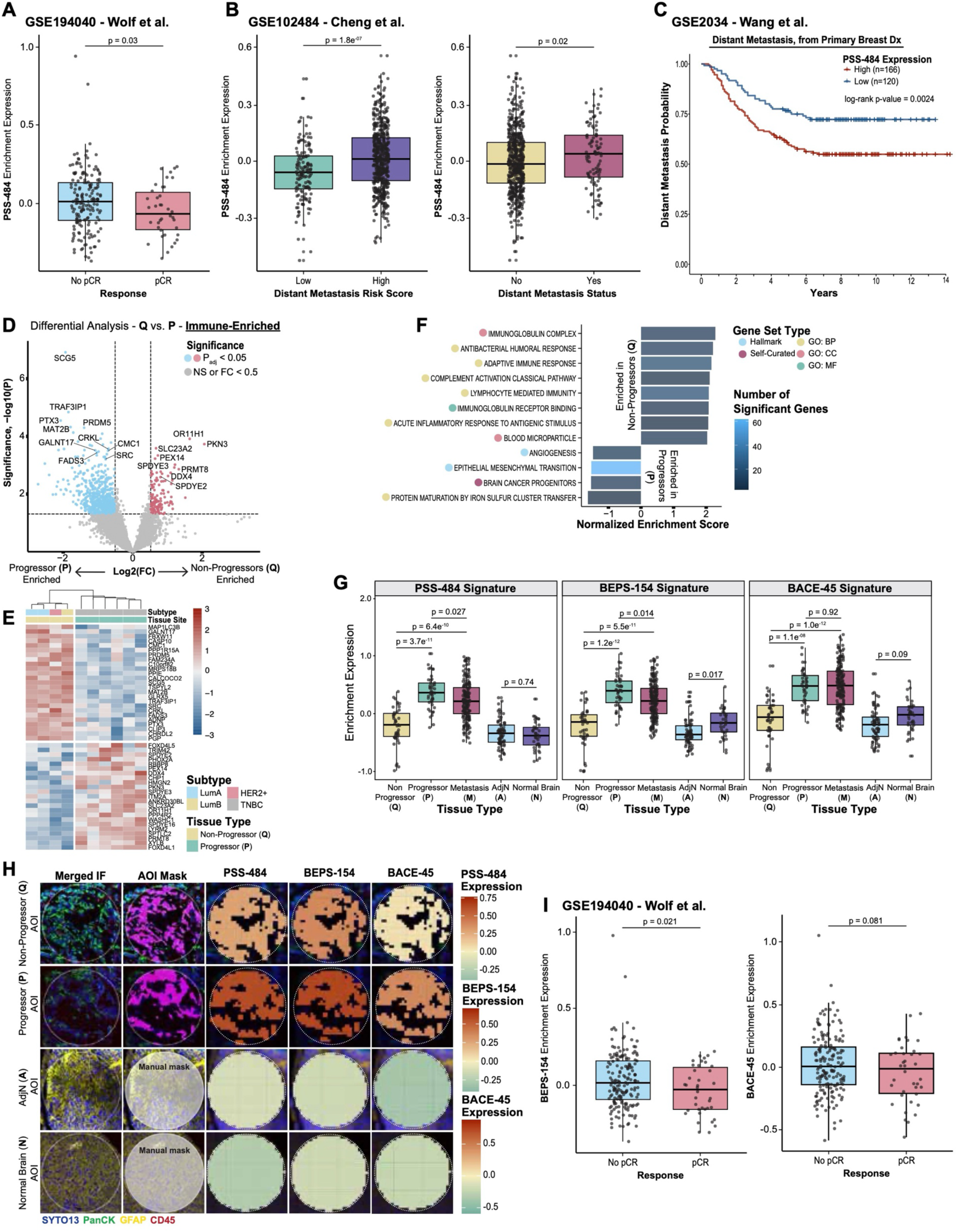
(Supportive Data to Main Figure 6). Validation of Progressor-Primary signatures and Differential expression analysis in Non-Progressor-Primary (Q) vs Progressor-Primary (P) Immune-Enriched AOIs. **(A)** Boxplot of PSS-484 enrichment expression by pathological complete response (pCR). n = 172 and 38 patients that did not or did achieve pCR; respectively (data from ref. [43]). **(B)** Boxplot of PSS-484 enrichment expression by (*left*) distant metastasis risk score and (*right*) distant metastasis status. n = 146 and 537 patients with Low and High risks scores; respectively. n = 582 and 101 patients who did not or did develop distant metastases; respectively (data from ref. [44]). **(C)** Kaplan-Meier curves of distant metastasis free survival (from date of primary breast cancer diagnosis) as a function of PSS-484 expression. Patients were segregated into two groups (High, n = 166; Low, n = 120) based on the enrichment expression of the PSS-484 signature in primary breast tumors (data from ref. [45]). The log-rank p-value was derived from comparing discretized predictors (high versus low PSS-484 expression). **(D)** Volcano plot of differentially expressed genes between Non-Progressor-Primary (n = 14) and Progressor-Primary (n = 10) Immune-Enriched AOIs. P-values were calculated using a mixed-effects model using chi-square (χ2) tests (two-sided) with p < 0.05 considered as significant. Genes with p > 0.05 and/or Log_2_(Fold Change) < 0.5 are considered not significant. **(E)** Heatmap of top 25 differentially expressed genes in Non-Progressor-Primary and Progressor-Primary Immune-Enriched AOIs. Tissue type and molecular subtype are indicated by colored bars. Heatmaps colored from blue to red according to Log_2_(FC) value of –3 to 3. **(F)** Gene set enrichment analysis of genes upregulated in Non-Progressor-Primary compared to Progressor-Primary Immune-Enriched AOIs. Gene set type and number of significant (p < 0.05) genes are denoted. **(G)** Boxplot of PSS-484, BEPS-154, and BACE-45 enrichment expression by tissue type. n = 43, 44, 182, 73, and 38 AOIs in Non-Progressor, Progressor, Metastasis, Adjacent Normal, Normal Brain; respectively. P values are based on nonparametric test (Kruskal– Wallis) followed by Dunn test for pairwise comparisons. **(H)** Representative spatial omics overlay of PSS-484, BEPS-154, and BACE-45 enrichment expression by tissue type. Pseudocolor merged IF image, AOI masked image, and signature enrichment overlay are shown. Signature enrichment overlays were generated using the SpatialOmicsOverlay R package (see Methods). **(I)** Boxplot of (*left*) BEPS-154 and (*right*) BACE-45 enrichment expression by pathological complete response (pCR). n = 172 and 38 patients that did not or did achieve pCR; respectively (data from ref. [43]). For boxplots in **A**, **B**, **G**, and **I**, middle line denotes the median, box edges indicate the 25th and 75th percentiles, and whiskers extend to the most extreme points that do not exceed ±1.5 times the interquartile range (IQR). P-values are based on paired Wilcoxon rank sum test. FC: fold change. GO: Gene ontology. BP: biological process. CC: cellular component. MF: molecular function. IF: immunofluorescence.

**Supplemental Figure 7.**
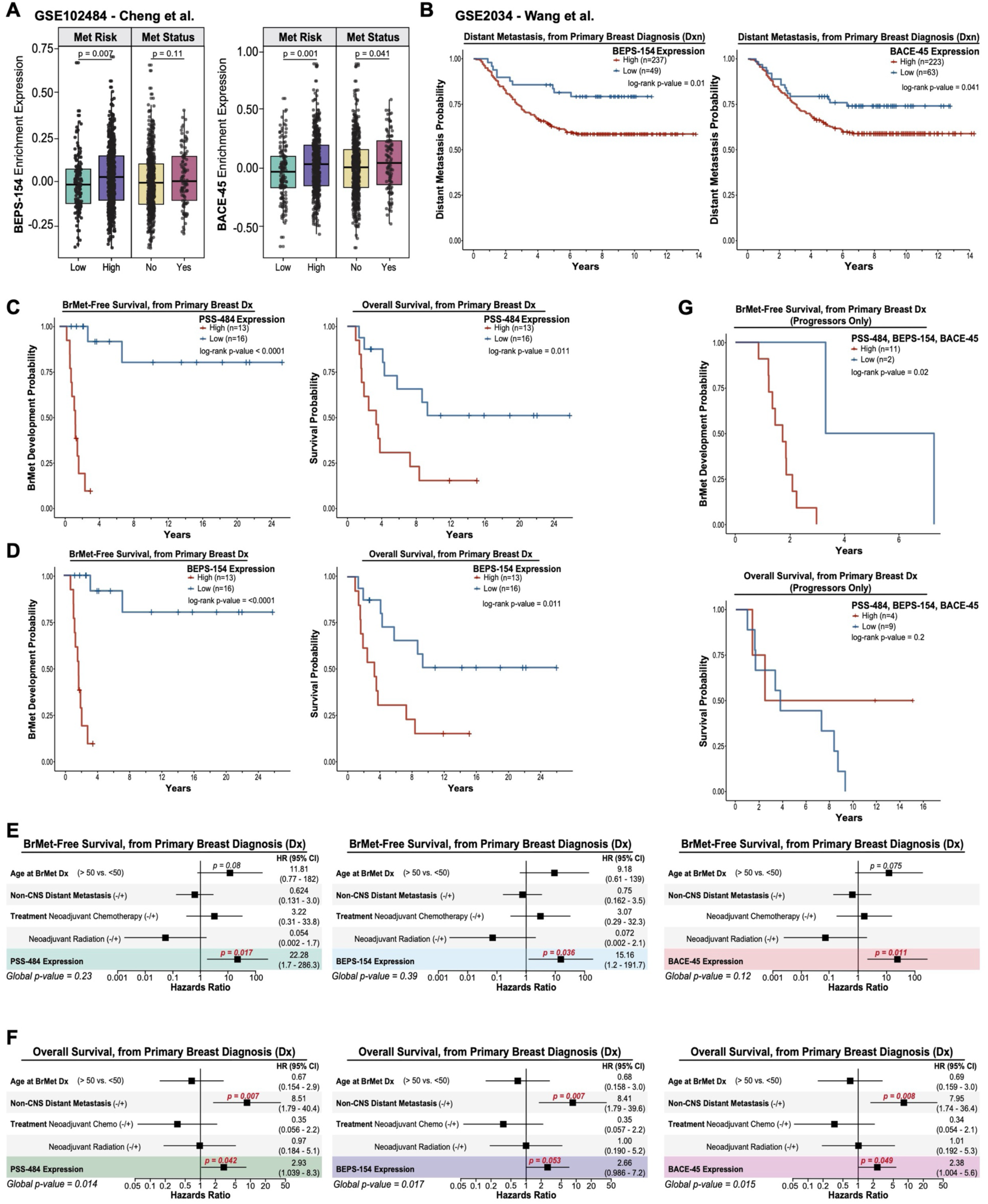
(Supportive Data to Main Figure 6). Validation of Progressor-Primary signatures and Progressor-Primary signatures impact on BMFS and OS. **(A)** Boxplot of BEPS-154 and BACE-45 enrichment expression by (*left*) distant metastasis risk score and (*right*) distant metastasis status. n = 146 and 537 patients with Low and High risks scores; respectively. n = 582 and 101 patients who did not or did develop distant metastases; respectively (data from ref. [44]). For boxplots in **A**, middle line denotes the median, box edges indicate the 25th and 75th percentiles, and whiskers extend to the most extreme points that do not exceed ±1.5 times the interquartile range (IQR). P-values are based on paired Wilcoxon rank sum test. **(B)** Kaplan-Meier curves of distant metastasis free survival (from date of primary breast cancer diagnosis) as a function of (*left*) BEPS-154 and (*right*) BACE-45 expression. Patients were segregated into two groups (BEPS-154: High, n = 237; Low, n = 49. BACE-45: High, n = 223; Low, n = 63) based on the enrichment expression of the BEPS-154 and BACE-45 signatures in primary breast tumors (data from ref. [45]). **(C)** Kaplan-Meier curves of *(left)* brain metastasis-free survival (BMFS) (from date of primary breast cancer diagnosis) and *(right)* overall survival (OS) (from date of primary breast cancer diagnosis) and as a function of PSS-484 expression. Patients were segregated into two groups for BMFS (High, n = 13; Low, n = 16) and OS (High, n = 13, Low, n = 16) based on the enrichment expression of the PSS-484 signature in Malignant AOIs from primary breast tumors. **(D)** Kaplan-Meier curves of *(left)* brain metastasis-free survival (BMFS) (from date of primary breast cancer diagnosis) and *(right)* overall survival (OS) (from date of primary breast cancer diagnosis) and as a function of BEPS-154 expression. Patients were segregated into two groups for BMFS (High, n = 13; Low, n = 16) and OS (High, n = 13, Low, n = 16) based on the enrichment expression of the BEPS-154 signature in Malignant AOIs from primary breast tumors. **(E)** Multivariable Cox proportional hazards regression analyses assessing the impact of clinical covariates and the expression of PSS-484 (*left*), BEPS-154 (*middle*), and BACE-45 (*right*) on brain metastasis-free survival (BMFS) (from date of primary breast cancer diagnosis). **(F)** Multivariable Cox proportional hazards regression analyses assessing the impact of clinical covariates and the expression of PSS-484 (*left*), BEPS-154 (*middle*), and BACE-45 (*right*) on overall survival (OS) (from date of primary breast cancer diagnosis). **(G)** Kaplan-Meier curves of *(top)* brain metastasis-free survival (BMFS) (from date of primary breast cancer diagnosis) and *(bottom)* overall survival (OS) (from date of primary breast cancer diagnosis) and as a function of PSS-484, BEPS-154, and BACE-45 expression in Progressor-Primary patients only. Patients were segregated into two groups for BMFS (High, n = 11; Low, n = 2) and OS (High, n = 4, Low, n = 9) based on the enrichment expression of the PSS-484, BEPS-154, and BACE-45 signatures in Malignant AOIs from primary breast tumors. (Note: Kaplan-Meier curves for all 3 signatures for BMFS were identical, so they are represented as one plot. Kaplan-Meier curves for all 3 signatures for OS were identical, so they are represented as one plot.) In **B**, **C**, **D**, and **G**, the log-rank p-value was derived from comparing discretized predictors (high versus low expression of PSS-484, BEPS-154, or BACE-45). In **E** and **F**, for each covariate, the hazard ratio (HR) and its 95% confidence interval (CI), denoted by bars, are presented. In **E** and **F**, p-values for each covariate were calculated using the Wald statistic. p < 0.05 are significant and denoted in bold red text. p-values near significance are also noted. Global p-values were calculated using the log-rank statistic.

**Supplemental Figure 8.**
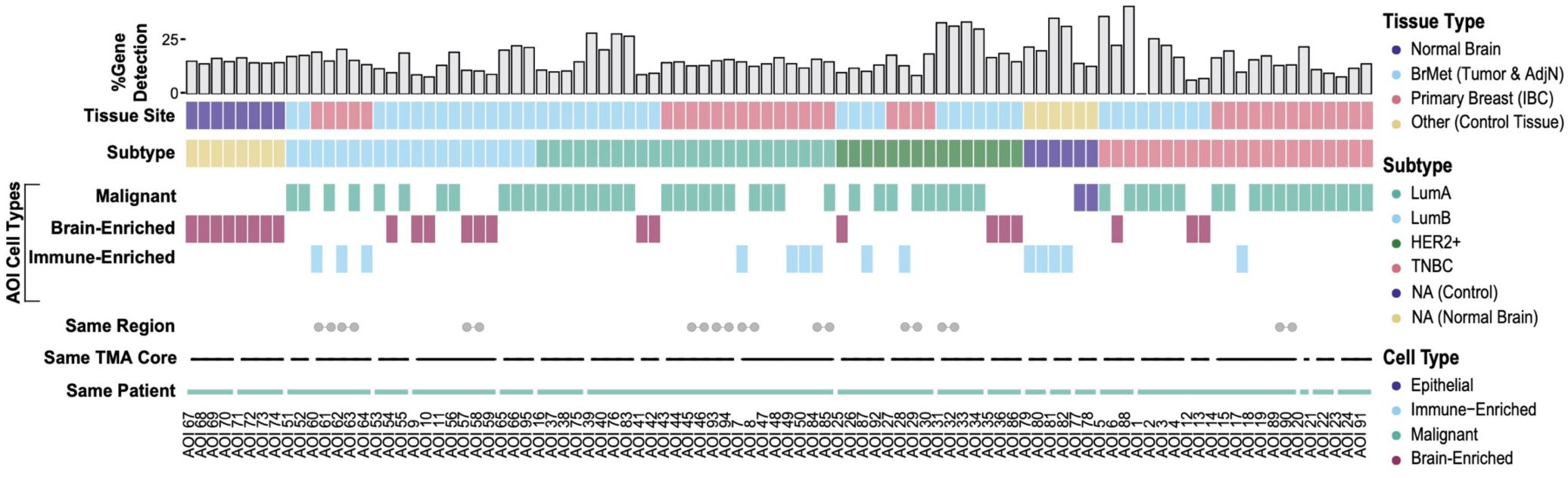
(Supportive Data to Main Figure 1). Representative summary of AOI annotations from study cohort. Each column represents 1 of 96 AOIs, detailing metadata on patient ID, TMA core, region, cell type, tissue type, molecular subtype, and % gene detection based on QC metrics (Methods).

**Supplemental Figure 9.**
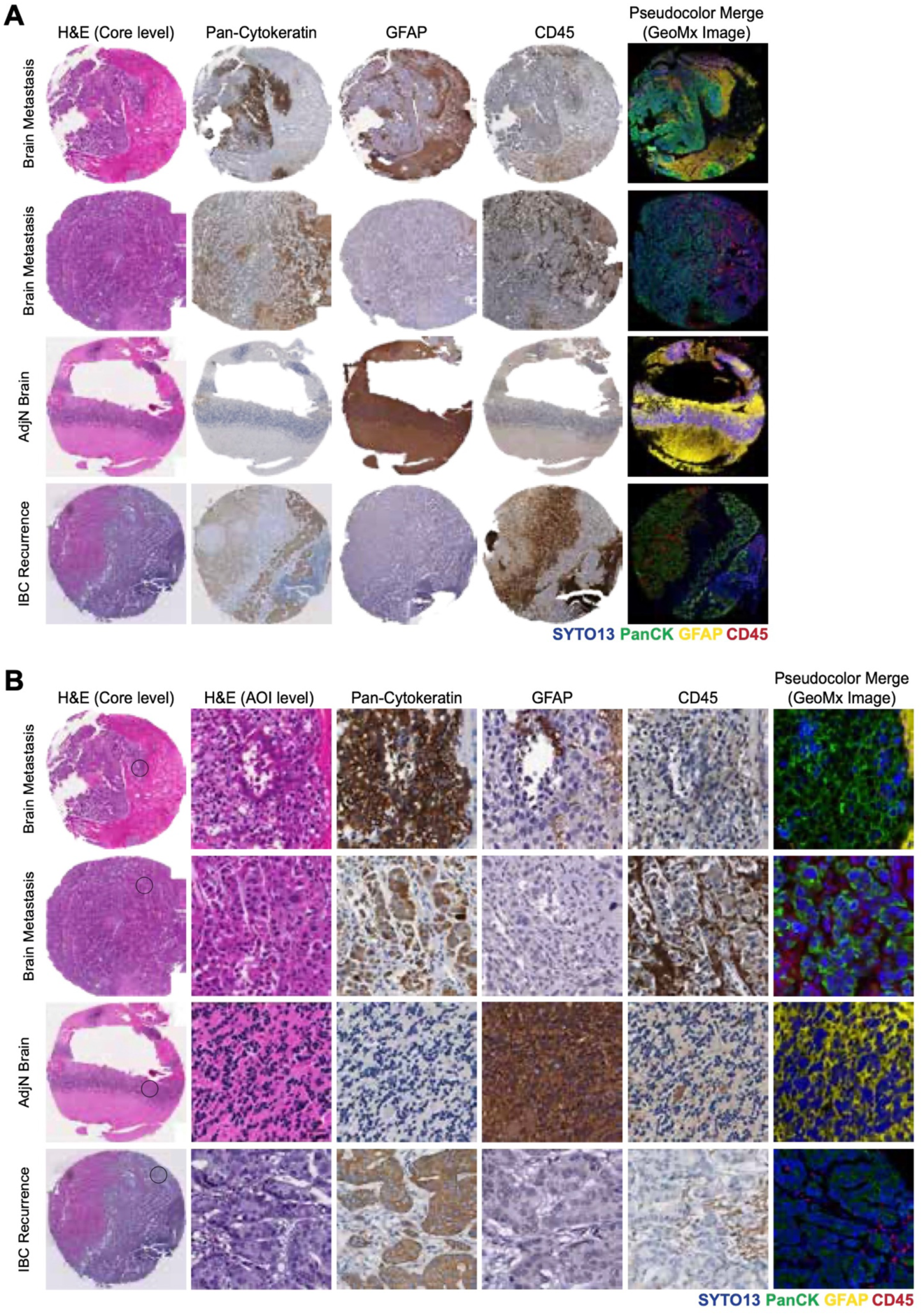
(Supportive Data to Main Figure 1 and Supplemental Table 1). Validation of GeoMx Imaging. **(A-B)** Representative images of **(A)** whole tissue microarray cores and **(B)** regions with cores (indicated by black circle). H&E stain, IHC of Pan-Cytokeratin, GFAP, and CD45 antibodies, and pseudocolor merged IF (from GeoMx DSP instrument) images are shown.

**Supplemental Figure 10.**
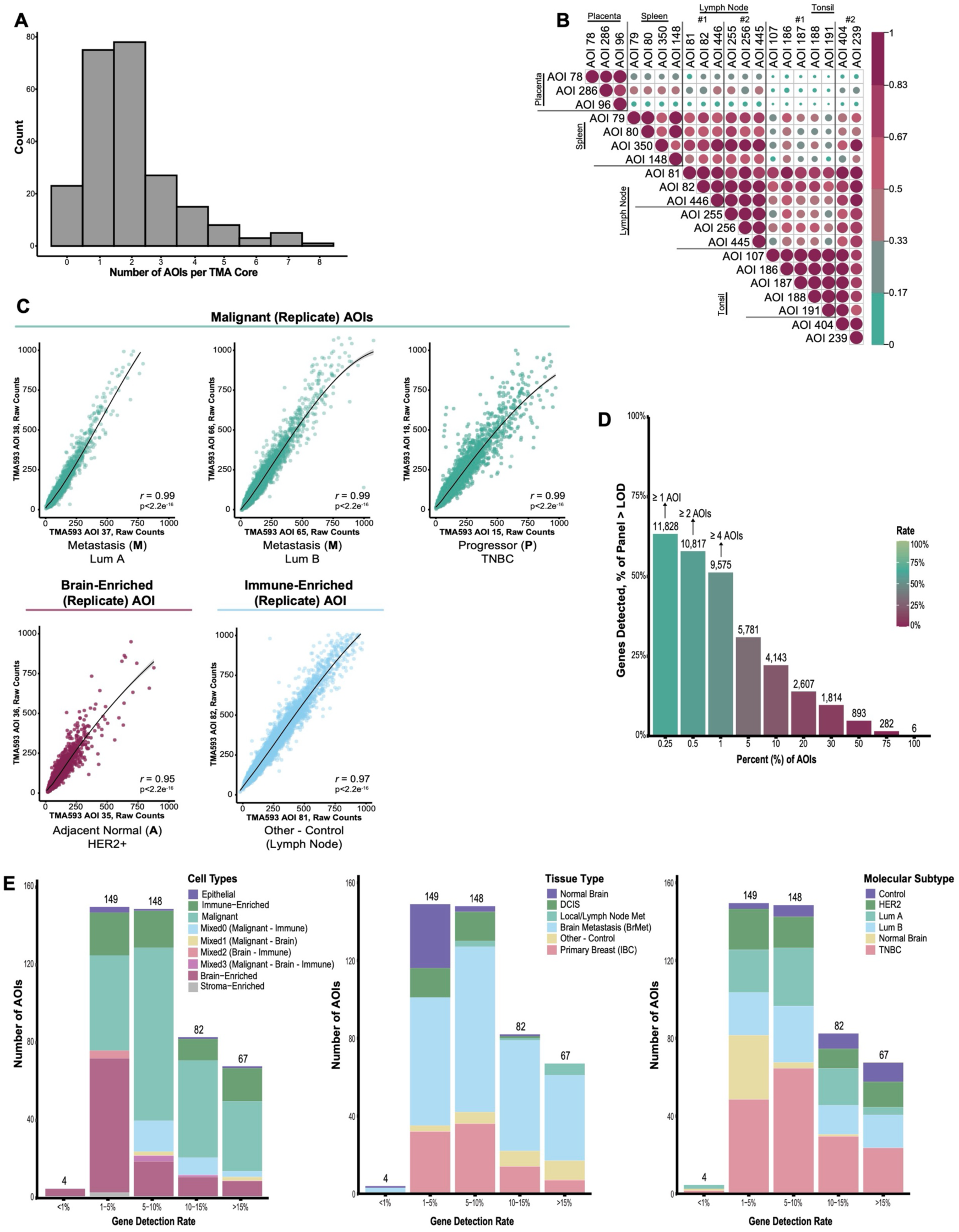
(Supportive Data to Main Figure 2 and Supplemental Table 4 & 5). Intra– and Inter-AOI variations and DSP gene detection levels. **(A)** Histogram representing the distribution of the number of AOIs per individual TMA core. **(B)** Correlation matrix heatmap showing the pairwise correlation coefficients (Pearson) between Control AOIs using Q3 normalized counts (n = 9575 genes). Heatmap colored from green to red according to correlation coefficient score 0.0 to 1.0. n = 3, 4, 6, and 7 AOIs from placenta, spleen, lymph node, and tonsil tissues; respectively. Control tissues were present across all three TMA slides (TA593, TA594, TA595). **(C)** Representative scatterplots depicting the relationships between raw genes counts (n = 18677 genes) observed in two replicate AOIs from an individual TMA core. Replicate AOIs from Malignant, Brain-Enriched and Immune-Enriched AOI types are shown. Each point represents a single gene. A regression line (loess smoothing method) is fitted to the data. The correlation coefficient (Pearson, r) and p-value are provided for each plot. **(D)** Barplot depicting the total number of genes above the Limit of Detection (LOD) in specific percentages of AOIs. Y-axis indicates the gene count as a percentage of the whole transcriptome panel (n = 18677 genes). Given the biological diversity of this dataset, 1% of AOIs was utilized as a cutoff (Methods). Genes not detected in at least 1% of AOIs were excluded from downstream analysis. **(E)** Stacked barplots showing the number of AOIs per range of gene detection rate. Plots are stratified by (*left*) cell type, (*middle*) tissue type, and (*right*) molecular subtype.

**Supplemental Figure 11.**
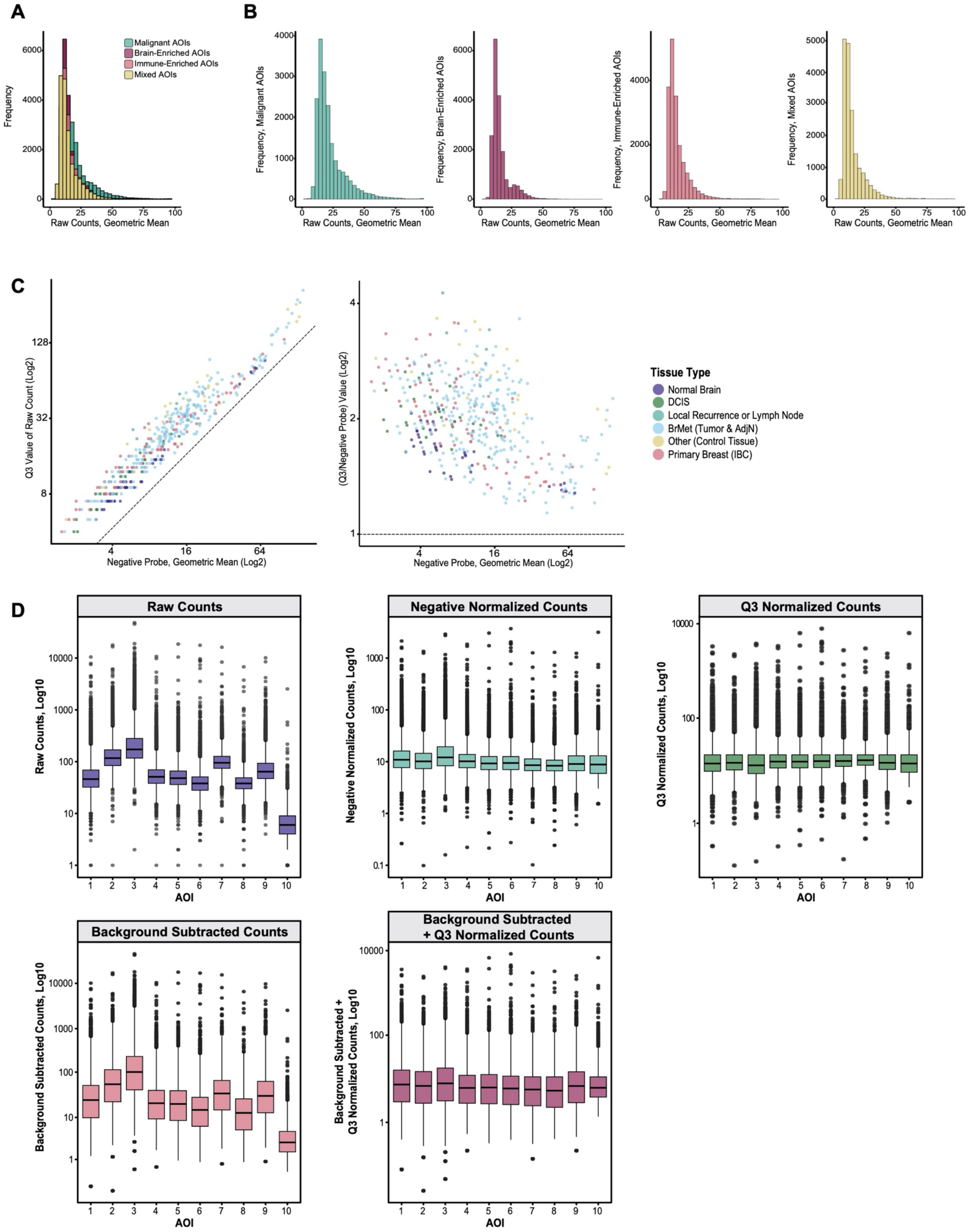
(Supportive Data to Main Figure 2). AOI raw counts and Normalization. **(A-B)** Histogram representing the distribution of raw genes counts (n = 18677 genes, geometric mean) overlaid (**A**) and plotted separately (**B**) for Malignant, Brain-Enriched, Immune-Enriched, and Mixed [Mixed0, Mixed1, Mixed2, Mixed3] AOIs. **(C)** Scatterplots depicting the relationship between Q3 normalized counts and the geometric mean of negative control probes. Each point represents an AOI (n = 450) and is annotated by tissue type. Dotted line indicates a 1:1 proportion of Q3 to negative probe counts. Scatterplot shows all AOIs are above this line, indicating a strong separation between Q3 counts and negative probe counts (Methods). **(D)** Representative boxplots of AOIs #1-10 before (raw counts) and after various normalization methods (negative probe normalization, Q3 normalization, background subtraction, and background subtraction followed by Q3 normalization). For boxplots in **D**, middle line denotes the median, box edges indicate the 25th and 75th percentiles, and whiskers extend to the most extreme points that do not exceed ±1.5 times the interquartile range (IQR); further outliers (minima and maxima) are marked individually as black points beyond the whiskers.

**Supplemental Figure 12.**
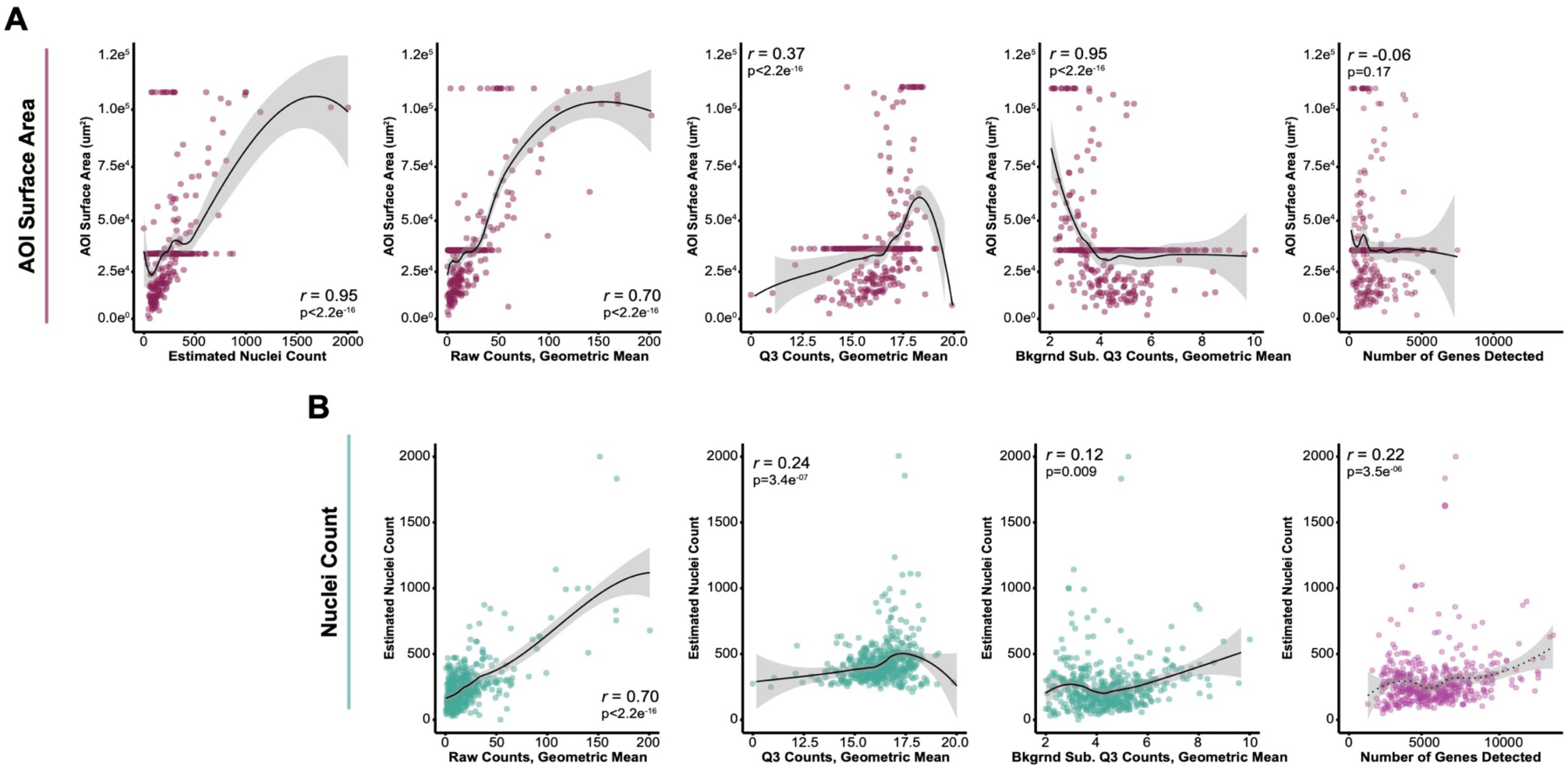
(Supportive Data to Main Figure 2). Correlations between AOI surface area, nuclei count, and gene counts. **(A)** Scatterplots depicting the relationships between AOI surface area and the geometric mean of: raw gene counts, Q3 normalized gene counts, background subtraction with Q3 normalization counts, and the number of genes detected. **(B)** Scatterplots depicting the relationships between AOI estimated nuclei counts and the geometric mean of: raw gene counts, Q3 normalized gene counts, background subtraction with Q3 normalization counts, and the number of genes detected. Each point represents an AOI (n = 450). For **A** and **B**, a regression line (loess smoothing method) is fitted to the data. The correlation coefficient (Pearson, r) and p-value are provided for each plot.

